# Dual AAV amelioration of Lama2-null muscular dystrophy and neuropathy

**DOI:** 10.64898/2026.02.09.704845

**Authors:** Karen K. McKee, Peter D. Yurchenco

## Abstract

The *dy^3K^/dy^3K^* Lama2**^-/-^** mouse is a model for the severe form of LAMA2-related dystrophy and peripheral neuropathy (LAMA2-RD). In the dystrophic mice, a compensating laminin subunit, Lmα4, that lacks polymerization and α-dystroglycan-binding activity, replaces the missing Lmα2 subunit. It was previously found that an α4-laminin can be modified with two small laminin-binding linker proteins, i.e. αLNNdΔG2’ and miniagrin to facilitate polymerization and α-dystroglycan binding respectively, to enable the key missing functions. Adeno-associated virus serotype 9 (AAV_9_) was used to deliver minigenes coding for the two proteins in dystrophic mice. AAV_9_-αLNNdΔG2’ utilized a universal CBh promoter while AAV_9_-miniagrin utilized either the CBh promoter or muscle-specific SPc5-12 promoter. The phenotype in the *dy^3K^/dy^3K^* mice was evaluated following i.v. postnatal injection with either AAV_9_ -αLNNdΔG2’ alone or in combination with AAV_9_- αLNNdΔG2’ + AAV_9-_ miniagrin. Double AAV treatment was found to substantially increase survival and ambulation, as well as increase forelimb grip-strength and improve muscle histology. Of note, the sciatic nerve amyelination characteristic of laminin α2-deficiency was prevented. While single treatment with αLNNdΔG2’ was inferior to double treatment for muscle strength and survival, it corrected the radial sorting deficit equally, revealing that enablement of laminin polymerization is a sufficient requirement for myelination.

**Highlights:**

- The *dy^3K^/dy^3K^* (Lama2^-/-^) mouse, a model for severe LAMA2-related dystrophy, expresses laminin-411 that is unable to polymerize or bind to α-dystroglycan (αDG).
- αLNNdΔG2’ and miniagrin are laminin-411-binding proteins that enable polymerization and αDG binding.
- AAV_9_ delivery of genes coding for αLNNdΔG2’ and miniagrin ameliorated the dystrophic phenotype in muscle and nerve (survival, growth, mobility, and grip-strength, muscle and nerve histopathology).
- Sciatic nerve amyelination was prevented by αLNNdΔG2’ alone.

## INTRODUCTION

Laminin α2-deficient <u>R</u>elated <u>D</u>ystrophy (*LAMA2-RD*) is an autosomal recessive disease that typically presents as a severe congenital muscular dystrophy (CMD) [1, 2] accompanied by peripheral neuropathy and brain anomalies. The prevalence varies from 1.79 per million in East Asians to 10.1 per million in Europeans [3]. Laminin-211 (a heterotrimer of α2, β1 and γ1 subunits, abbreviated Lm211 [4]) is the major laminin of skeletal muscle myofiber and peripheral nerve Schwann cell (SC) BMs. It is also found in blood-brain barrier (BBB) capillaries and glia limitans. Over 90% of *LAMA2* mutations result in a complete or substantial loss of protein subunit expression (*<u>M</u>uscular <u>D</u>ystrophy <u>C</u>ongenital type IA,* MDC1A)[5–7]. These LAMA2-RD patients express compensatory Lm411 in muscle BM [8]. Missense and in-frame deletion mutations, mostly mapping to the Lmα2 short-arm polymerization domain (LN) region, cause a milder ambulatory disease (≤5%) in which the peripheral neuropathy can be prominent [9–12]. The pathology in both variants consists of myofiber loss (largely through apoptosis), regeneration from satellite cell activation and proliferation, chronic inflammation and fibrosis accompanied by white matter brain hyperlucency detected by T2-weighted MRI, and peripheral nerve amyelination with reduced conduction [2]. Patients with the severe disease are so weak they never ambulate, develop severe joint contractures, and are at risk for dying of muscle wasting and, if untreated, respiratory failure in the first decade of life. Patients with defective α2-laminin polymerization present later in life with persistent weakness yet become ambulatory [11]. There is currently no cure for either [6].

The laminin α2-null mouse (*dy^3K^/dy^3K^*) is a model for the most severe manifestation of the human disease [13]. Successful treatment of the dystrophic mice is more challenging than the treatment of mice that only have a polymerization defect [14]. Notably, the *dy^3K^/dy^3K^* mice weigh only about half their normal wild-type (WT) counterpart at weaning, move sluggishly, and survive 8 weeks or less with half dying by 4 weeks of age [15].

Total muscle γ1-laminin concentrations in muscle extracts of these mice were found to be reduced by about three-fold compared to wild-type mice with laminin α4 as the principal replacement subunit [15]. It is has been found that Lm411 (α4β1γ1) has only weak adherence to BMs and easily dissociates from muscle BMs because it is unable to polymerize to create cooperatively of binding to receptors (through avidity rather than affinity) [16].

A characteristic of the different mouse models is the presence of a limb abnormality seen as a hindlimb retraction towards the body rather than outwards upon lifting the mouse by the tail. This is followed by gradual development of hindlimb paresis and contractures, prominent in *dy^2J^/dy^2J^* by 11 weeks, but mainly seen in *dy^3K^/dy^3K^* if survival extends beyond ∼6 weeks of age [15]. The hindlimb abnormality is associated with partial amyelination of the sciatic nerve and nerve roots, a late developmental defect of radial axonal sorting [17–19]. Important to this process are the interactions of α2- and α4-laminins with β1-integrins and α2-laminins with dystroglycan [17, 20, 21].

In the current study, we examined the effects of delivering the two laminin-binding “linker protein” genes to muscle and peripheral nerve by adeno-associated virus (AAV), an approach used in a variety of clinical trials to treat neuromuscular diseases [22–24]. The corresponding linker proteins enable laminin polymerization (αLNNdΔG2’) and increase cell anchorage through α-dystroglycan (miniagrin). Significant improvements were observed in both muscle and nerve.

## RESULTS

We employed AAV_9_ delivery of genes coding for the smaller αLNNdΔG2’ (compared to αLNNd) and miniagrin (*mag*) in *dy^3K^/dy^3K^*, comparing mice treated with AAV-CBh-αLNNdΔG2’ alone, or in combination with either universal CBh-*mag* or muscle-specific SPc5-12-*mag* (**Fig. 1**). Expression of both linker proteins was detected in muscle sections of doubly-treated (AAV_9_-CBh-αLNNd dG2’ + AAV_9_-CBh-*mag*) in sarcolemma, in Schwann cell endoneurium and perineurium, and (prominently for αLNNdΔG2’) microvessels (**Fig. 2**). Both proteins were detected at the expected intact size in immunoblots of muscle extracts.

**Fig. 1.**
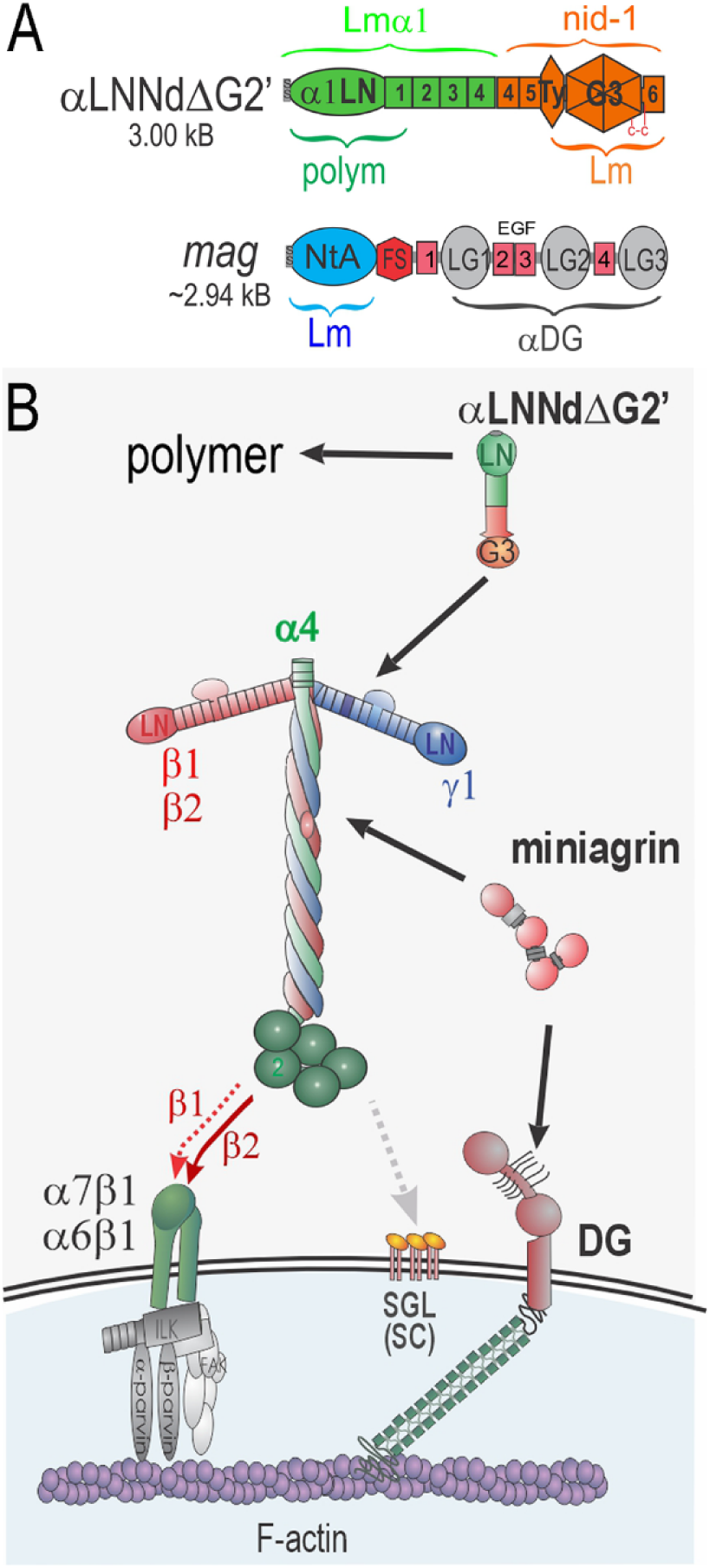
Linker proteins. **Panel A**. A shortened laminin-binding linker protein, αLNNdΔG2’, has an N-terminal polymerization domain (α1LN), tandem laminin-like EGF (LE) spacer domains, and a G3 domain complex (Ty thyroglobulin, G3 propeller and terminal EGF-6 domains) that binds to laminins at the nidogen-binding locus in the γ1 subunit. It enabled laminin polymerization when bound to a laminin that lacks either an αLN domain or the short arm containing the αLN domain. Lower construction. Miniagrin (*mag*) is an internally shortened version of muscle (non-neural) agrin and consists of a domain (NtA) that binds to the coiled-coil domain of laminins, connecting FS and EGF domains, and an LG/EGF complex that binds strongly to α-DG. It serves to anchor bound laminins to αDG and its linked cytoskeleton. **Panel B**. *Double linker protein strategy to modify* α*4-laminins so that they polymerize and anchor to* α*DG*. Lm411, a compensating neuromuscular laminin in the absence of Lm211, is unable to polymerize (lacks α4 short arm) or to bind to the αDG anchoring receptor. Lm411 and Lm421 attach to adhesive (but non-anchoring) sulfatides on Schwann cell surfaces. Lm421, unlike Lm411, binds to integrins comparable to other laminins but similarly is unable to bind to αDG. A combination of αLNNdΔG2’ and miniagrin corrects the polymerization and dystroglycan deficiencies.

**Fig. 2.**
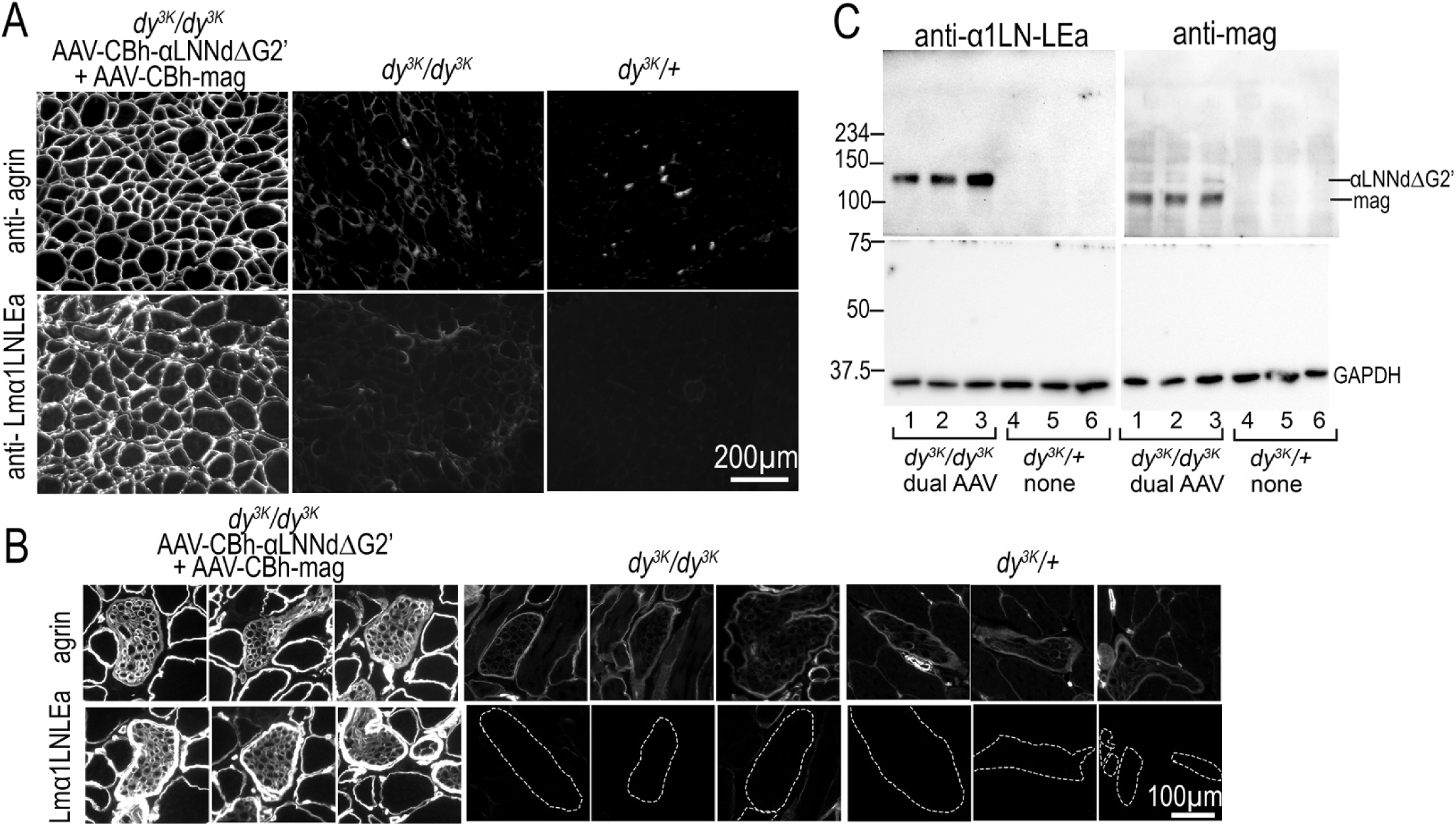
Expression of linker proteins in dy^3K^/dy^3K^ muscle and nerve. Dystrophic mouse pups were injected with AAV-CBh-αLNNd and AAV-*mag* at post-natal day 1 and sacrificed for muscle (triceps) immunostaining and tissue extraction at 6-weeks of age. **A** and **B**: Proteins miniagrin and αLNNdΔG2’ were detected in skeletal muscle (A) and nerve branches (B) in cross-sections with antibodies to agrin and Lmα1LN-LEa. Agrin immunofluorescence was substantially higher for miniagrin compared to endogenous agrin. Lmα1LN-LEa antibody detected αLNNdΔG2’ with untreated nerve outlines shown (borders determined from Lmα4 counterstains). **C**: Western immunoblots (SDS-PAGE, reduced) from 3 mice reveal intact protein bands for αLNNdΔG2’ and miniagrin at the expected molecular masses. Staining was evenly distributed in BMs across multiple tissue sections for both antibodies.

Mouse survival was increased more than two-fold with CBh-αLNNdΔG2’ alone and further increased with either of the two double AAV treatments. No spontaneous deaths were observed in the double-AAV-treated mice by 15 or 25 weeks (**Fig. 3A**), at which time the mice were sacrificed for tissue analysis. The weights of untreated *dy^3K^/dy^3K^* mice were substantially below that of controls and increased only slightly with age compared to controls. The weights of mice treated with only αLNNdΔG2’ were essentially the same as those of untreated mice. The weights of double AAV-treated *dy^3K^/dy^3K^* mice, on the other hand, regardless of whether *mag* expression was universal or muscle-specific, were increased to levels between untreated *dy^3K^/dy^3K^* and wildtype controls (**Fig. 3B**). The weights of these mice increased with advancing age, but with lower slopes compared to controls.

**Fig. 3.**
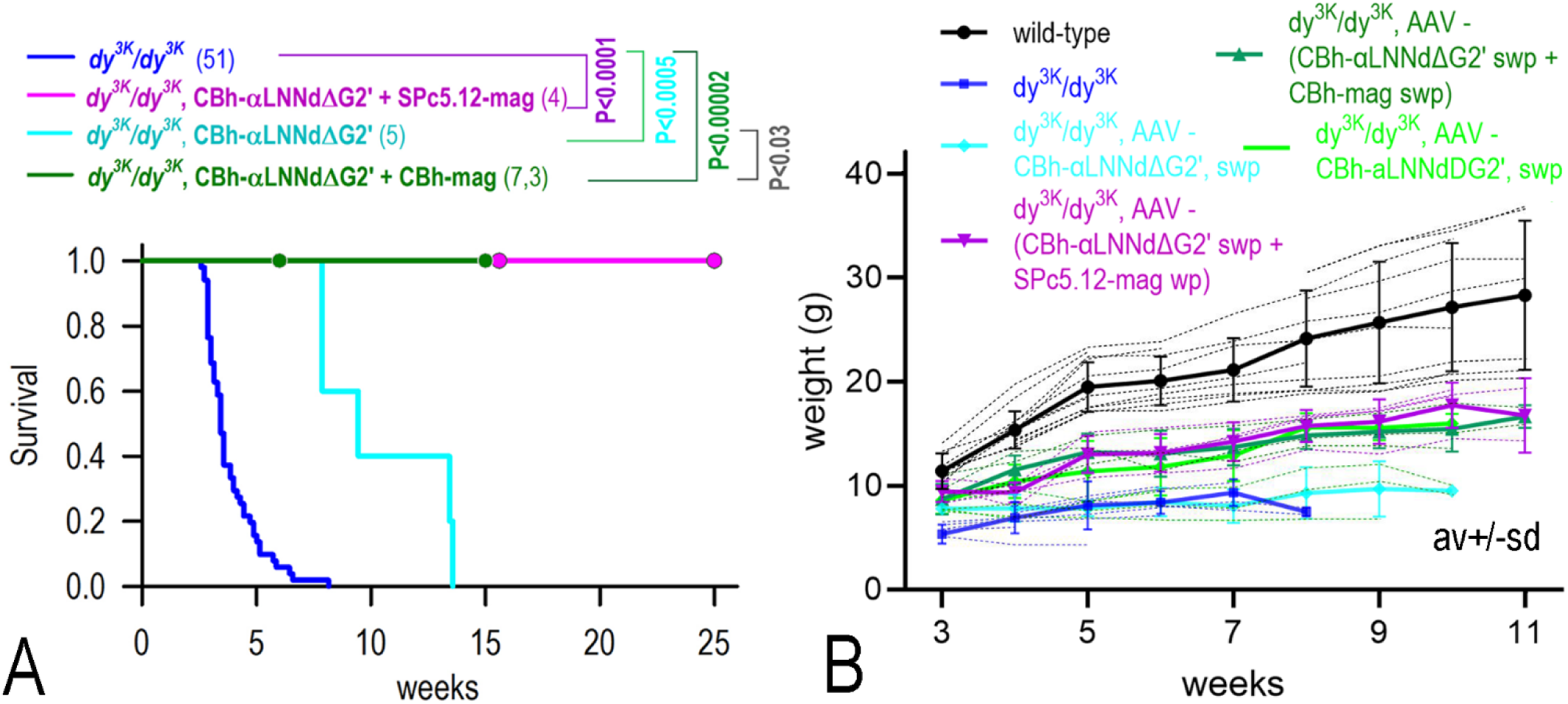
**A.** *Kaplan-Meier and Log-Rank Survival of dy^3K^/dy^3K^ mice*. Survival plots for dystrophic mice, untreated or treated with AAV-CBh-aLNNdΔG2’, AAV-CBh-αLNNdDG2’ + AAV-SPc5-12-*mag*, and AAV-CBh-αLNNdΔG2’ + AAV-SPc5-12-*mag*. Open circles indicate sacrifice for tissue analysis (pairwise Holm-Sidak comparisons; number of mice indicated in brackets). **B**. *Weights of dy^3K^/dy^3K^ mice*. Av ± s.d. and single values (thin dashed lines) shown. Untreated *dy^3K^/dy^3K^* mice were smaller and substantially lighter than wild-type mice. Double AAV treatments increased weights unlike AAV-CBh-αLNNdΔG2’ alone*. AAV doses*: CBh-αLNNdΔG2’ at 2 x 10^11^ vg/gm; CBh-αLNNdΔG2’ + SPc5-12-*mag* at 1.4 x 10^11^ vg/gm + 3.8 x 10^11^ vg/gm; CBh-αLNNdΔG2’ + CBh-*mag*, each at 2.8 x 10^11^ vg/gm.

Forelimb specific grip-strengths (grams force/grams weight) were measured from 3 to 11 weeks of age (**Fig. 4**). Treatment of *dy^3K^/dy^3K^* with AAV-CBh-αLNNdΔG2’ alone yielded a small (but not significant) increase over no treatment, while treatments with AAV-(CBh-αLNNdΔG2’ + SPc5-12-mag) or AAV-(CBh-αLNNdΔG2’ + CBh-mag) yielded greater (significant) improvements of grip strength. Both double AAV treatments increased forelimb grip-strength compared to untreated *dy^3K^/dy^3K^* with double CBh treatment exhibiting longer gripstrength persistence with increasing age. The dystrophic mice expressing αLNNdΔG2’ (with or without *mag*) did not develop the characteristic hindlimb phenotype seen upon lifting by the tail, regardless of age examined.

**Fig. 4.**
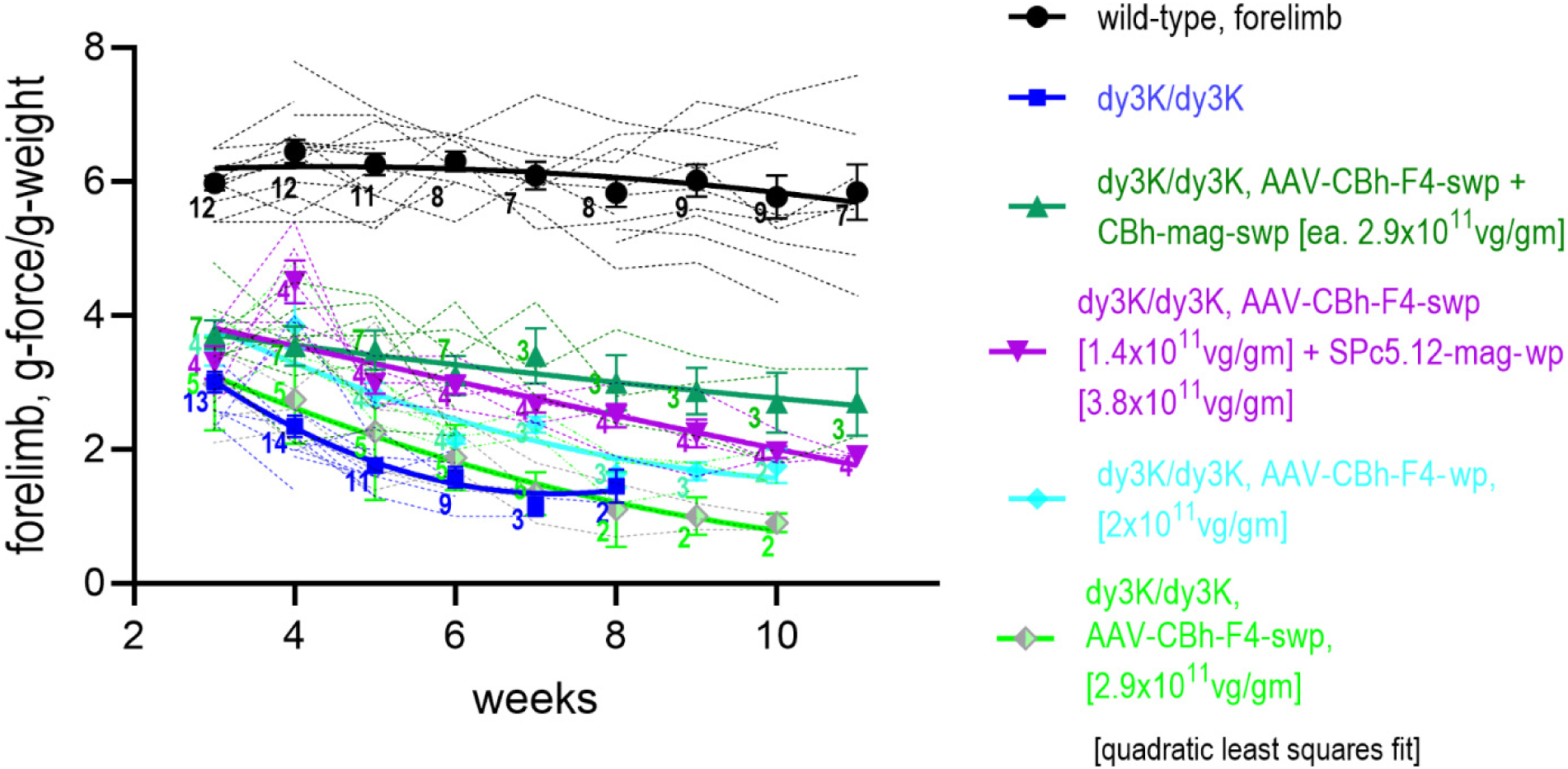
Forelimb grip-strength of dy3K mice. Specific grip-strengths are shown for the conditions listed in the legend. Av ± s.e.m. and individual plots (thin dashed lines) shown. Number of mice shown adjacent to each average data point. Pairwise comparisons (Tukey’s test) of a 2-way ANOVA of mean values revealed differences between *dy^3K^/+* and all other conditions (P<0.001), and between untreated and AAV-treated *dy^3K^/dy^3K^* with CBh-αLNNdΔG2’ + CBh-*mag* (P<0.002) and AAV-CBh-αLNNdΔG2’ + SPc5-12-*mag* (P=0.04). Other comparisons were not significant.

Mouse ambulation, examined by video recording steps, shown at 1 sec. intervals (**Fig. 5**), illustrated the difference in ambulatory movement between untreated and treated dystrophic mice. Untreated *dy^3K^/dy^3K^* mice typically remained stationary (as shown) with occasional ambulation. Dystrophic mice expressing only αLNNdΔG2’ showed considerably improved mobility as did dystrophic mice treated with αLNNdΔG2’ + miniagrin. Longer 10 min. duration recordings at 6, 11, and 15 weeks, measuring horizontal orthogonal light-beam interruptions, are shown in **Fig. 6**. Increased ambulation relative to untreated *dy^3K^/dy^3K^* was seen with all AAV treatments at 6 weeks. After that, ambulation (that could now only be compared to wild-type animals) remained elevated, but still never as high as seen in wild-type animals. Three video recordings showing ambulation are provided in the **Supplement** for *dy^3K^/dy^3K^* alone at 6 weeks (**Video-1**), *dy^3K^/dy^3K^*, AAV-CBh-αLNNdΔG2’ at 10 weeks (**Video-2**) and *dy^3K^/dy^3K^*. AAV-CBh-αLNNdΔG2’ + CBh-mag at 9.5 weeks (**Video-3**).

**Fig. 5.**
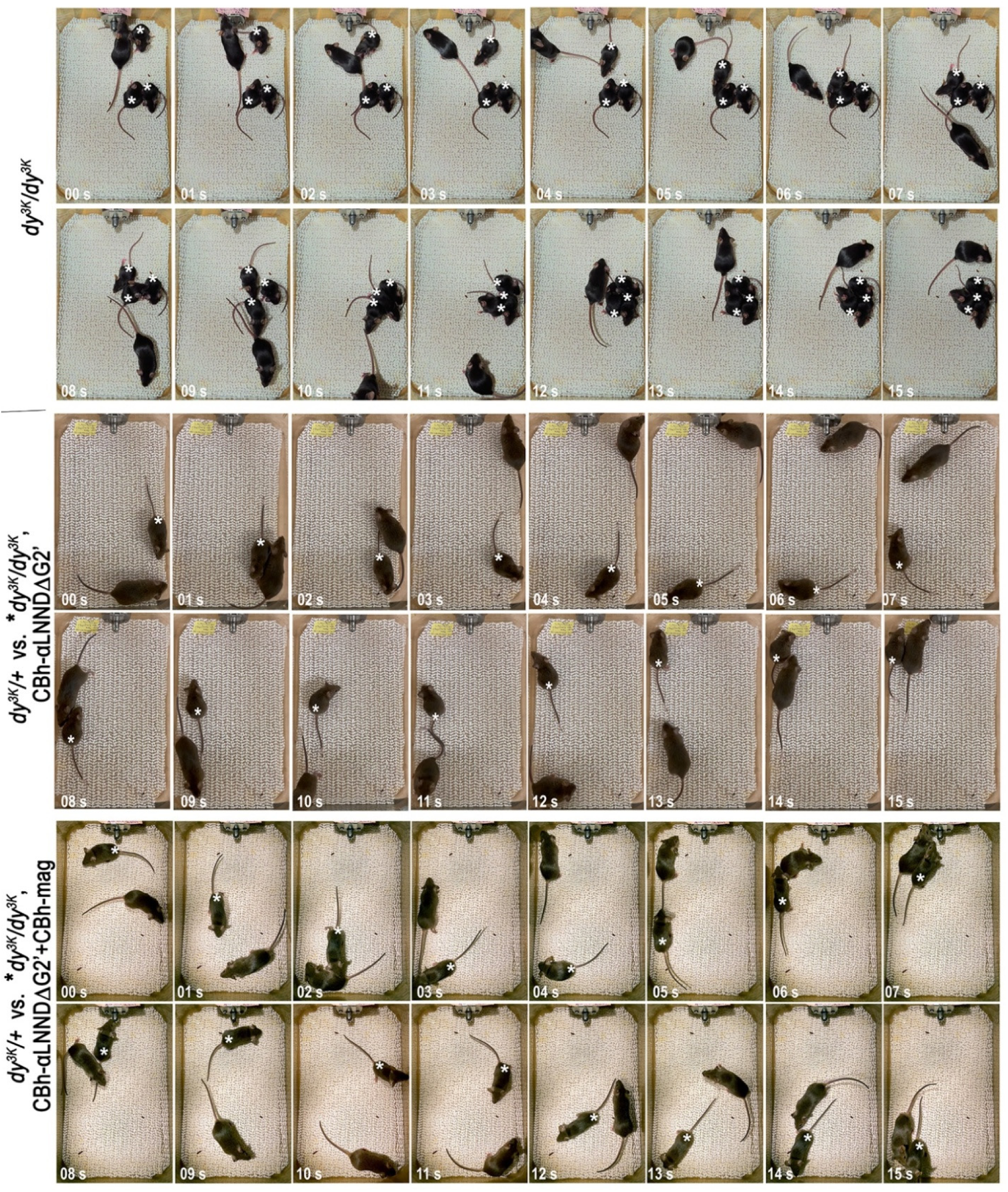
*Ambulation of (***A)** *a dy^3K^/+* with *3 dy^3K^/dy^3K^* mice at 6 weeks., (**B)** *a dy^3K^/+* mouse with a *dy^3K^/dy^3K^* treated with AAV-CBh-αLNNdΔG2 at 10 weeks., and (**C**) a *dy^3K^/+* mouse with a *dy^3K^/dy^3K^* treated with AAVCBh-αLNNdΔG2’ + CBh-mag at 9-1/2 wks. Mice were placed into a new cage environment. Representative one-second interval stills are shown over a 15 sec period. The untreated *dy^3K^/dy^3K^* mice remained nearly stationary (marked with asterisks) while the single and double AAV treated mouse (white asterisks) walked at increased levels.

**Fig. 6.**
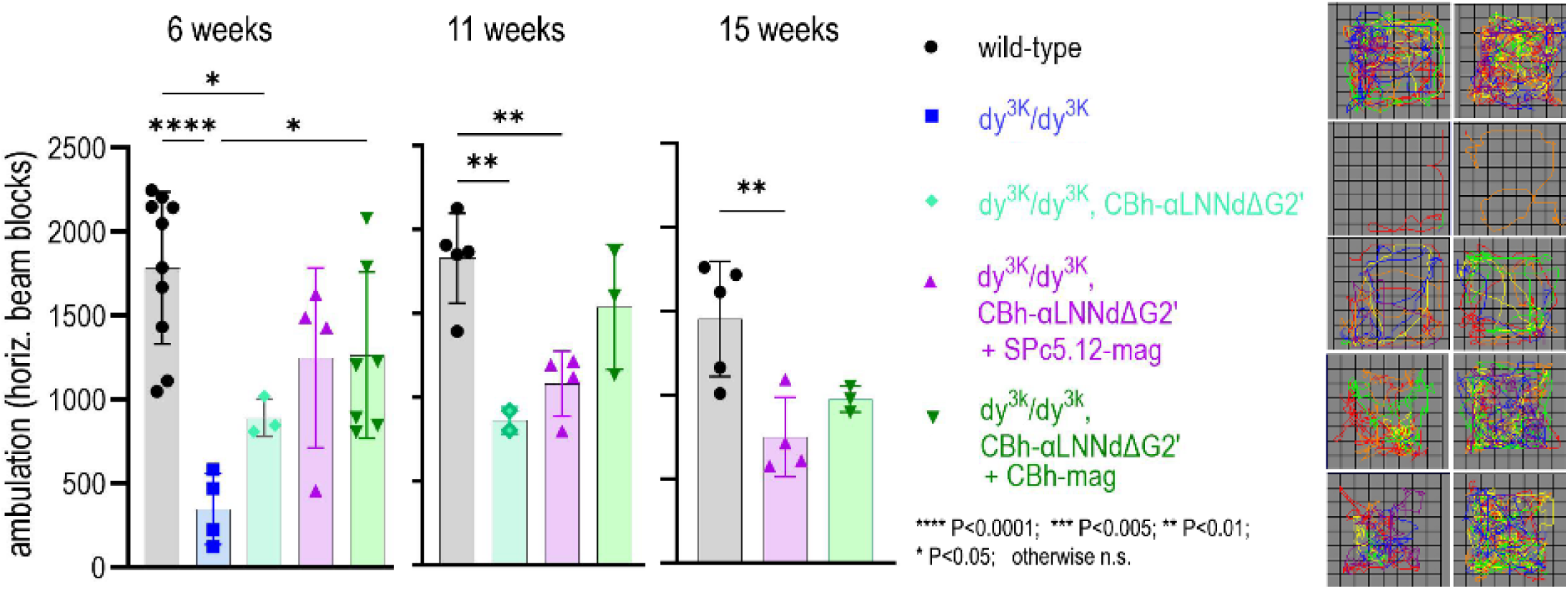
*Relative 10 min. walking distances of WT (dy^3K^/+), dy^3K^/dy^3K^ and dy^3K^/dy^3K^ treated with either AAV-CBh-αLNNdΔG2’, AAV-CBh-αfLNNdΔG2’ + SPc5-12-mag, or AAV-CBh-αLNNdΔG2’ + AAV-mag.* Av ± s.d. with individual mouse values shown [**** P<0.0001; *** P<0.001; ** P<0.01; * P<0.05, 1-way Anova with Tukey’s multiple comparisons test]. Path tracing 6-wk. examples shown at right. Untreated *dy^3K^/dy^3K^* exhibited little activity (and did not survive to be recorded at later times). An intermediate increase in activity was seen with the single AAV. Double AAV treatment resulted in further ambulatory activity.

Muscle histology was evaluated at 6 weeks of age (when untreated *dy^3K^/dy^3K^* mice were still available for comparison) comparing wild-type (*dy^3K^/dy^3K^* or +/+) and *dy^3K^/dy^3K^* with double AAV-CBh linker treatment. Examination of triceps (**Fig. 7**), rectus femoris (**Supplement Fig. 1**), plantaris (**Supplement Fig. 2a**) and small muscles of the distal forelimb (**Supplement Fig. 3**) revealed that the untreated *dy^3K^/dy^3K^* muscles were small compared to wild-type with greater variation of size, loss of angulation, and increased in central nucleation, a characteristic of regenerating muscle. The dystrophic mice treated with double CBh-AAV (αLNNdΔDG2’ and *mag*) were larger with an increased number of myofibers and decrease in fibrosis as determined from the ratio of collagen-stained area/total area. Importantly, there was almost a halving of the fraction of central nuclei, indirect evidence for reduction of regeneration following myofiber death. In general, larger muscles were less severely affected by the dystrophy and more responsive to linker protein expression compared to small forelimb muscles involved in gripping. Hindlimb muscles were also evaluated in adults at 10-15 weeks, with images shown for plantaris (**Suppl. Fig. 2b**). These revealed similar morphological relationships with wild-type controls as seen at earlier ages.

**Fig. 7.**
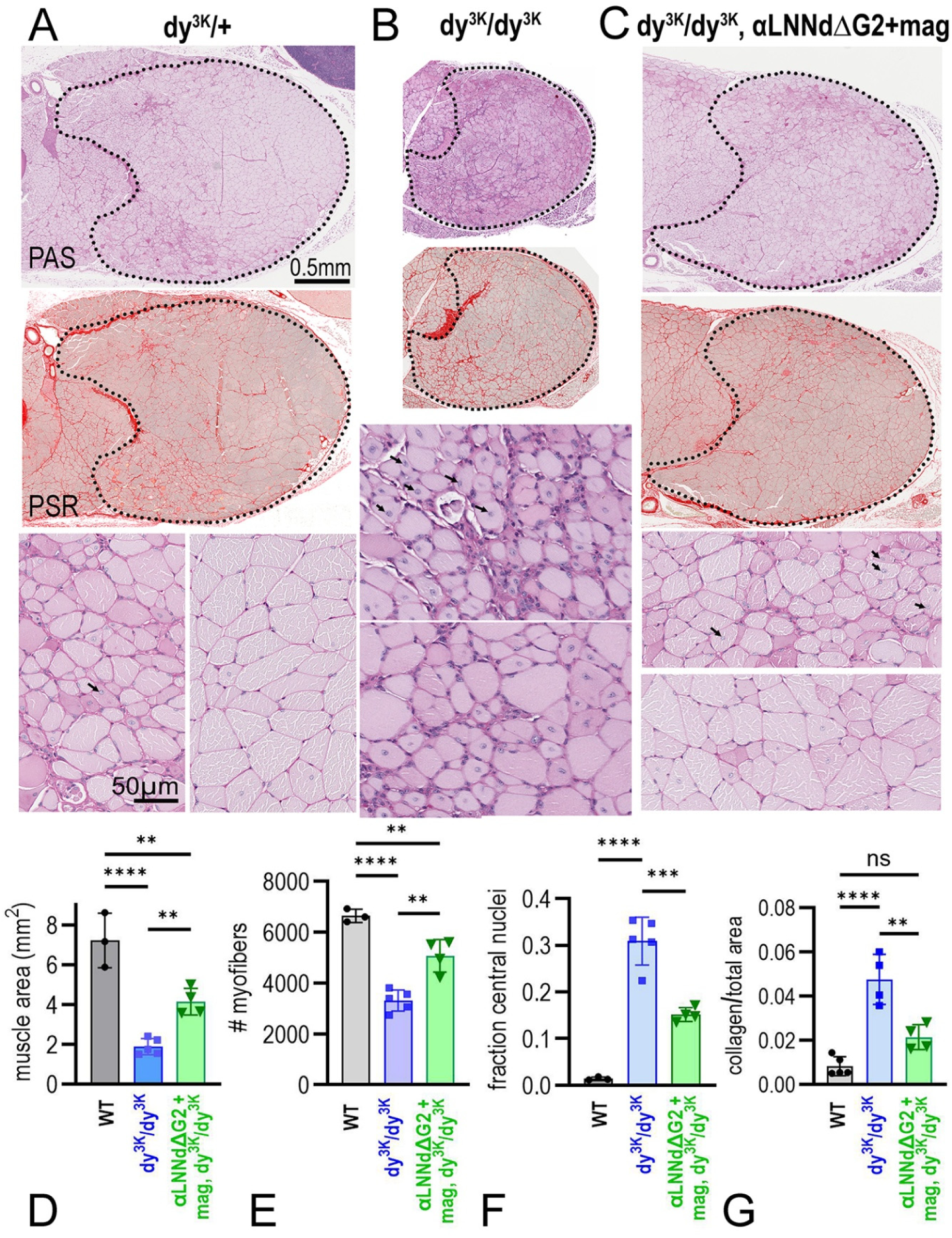
Histopathology of triceps muscle. Forelimb muscle from 6-week-old wild-type, untreated *dy^3K^/dy^3K^*, and *dy^3K^/dy^3K^* treated with AAV(CBh-αLNNdΔG2’ + AAV-CBh-*mag*) mice were stained with PAS and PSR. [**** P<0.0001; *** P<0.001; ** P<0.01; * P<0.05, 1-way Anova with Tukey’s multiple comparisons test]. Muscle areas and cross-sectional myofiber counts were increased while the fraction of central nuclei (regenerating fibers) and collagenous areas were decreased in the double-AAV treated *dy^3K^/dy^3K^* muscle.

Immunostaining of triceps from 6-week-old mice (**Supplement Fig. 4**) for laminin β1, β2 and γ1 subunits revealed reductions of these subunits in muscle sarcolemma and in small brachial nerve branch endoneurium compared to wild-type. Laminin α4 staining in muscle changed from a bright microvascular pattern to one that was present in muscle sarcolemma and reduced in microvasculature in *dy^3K^/dy^3K^*. Small brachial plexus-branch nerve endoneurium and epineurium were reduced for the above subunits in untreated *dy^3K^/dy^3K^* with little change or small increases following double AAV treatment. For muscle sarcolemma, the intensity of staining changes in *dy^3K^/dy^3K^* following AAV treatment were similar to those observed in *dy^3K^/dy^3K^* with transgenic expression of αLNNd plus miniagrin [15]. Laminin α5 staining was increased in sarcolemma of untreated *dy^3K^/dy^3K^* and following AAV treatment (**Supplement Fig 5**). However, we note a previous study revealed that this subunit represents a minor fraction in muscle [15]. α-Dystroglycan, present in wild-type sarcolemma, was increased in untreated *dy^3K^/dy^3K^* and greatly increased in following treatment with αLNNdΔG2’ + miniagrin delivered with a universal promoter (much more so than seen in *dy^3K^/dy^3K^* expressing the αLNNd and mag transgenes [15]). Small intramuscular nerve branch α-dystroglycan, on the other hand, was strongly present in perineurium but barely detectable in endoneurium. This is not surprising given the specialized location of α-dystroglycan in small narrow disks in the nodes of Ranvier [25], unlikely to be detected in muscle cross-sections.

Immunostaining intensity of integrin β1 (**Supplement Fig. 6**) in wild-type muscle was present in sarcolemma and strong in capillaries. Perlecan was used as a counterstain because it changes little among the different conditions. The capillary intensity of integrin β1 in untreated *dy^3K^/dy^3K^* was substantially decreased in capillaries and sarcolemma. BMs (especially those of capillaries) were increased in *dy^3K^/dy^3K^* expressing the two linker proteins. Integrin α7 sacrolemmal and capillary staining was substantially decreased in untreated *dy^3K^/dy^3K^* and greatly increased in *dy^3K^/dy^3K^* expressing the two linker proteins. Endoneurial integrins β1 and α7 were present at low intensity in *dy^3K^/+* (surrounded by bright perineurium), decreased in untreated *dy^3K^/dy^3K^*, and increased in double linker protein treated *dy^3K^/dy^3K^*.

The number of inflammatory macrophages (F4/80 staining), absent in wild-type and present in untreated *dy^3K^/dy^3K^* muscle, was considerably decreased in double linker protein *dy^3K^/dy^3K^* treated muscle (**Supplement Fig. 7**). This is similar to what was observed in *dy^3K^/dy^3K^* muscle without and with muscle-specific αLNNd and miniagrin double-transgene expression [15].

We examined sciatic nerve (**Fig. 8**) by light and electron microscopy. Multiple patches of naked axons (amyelination) of mixed caliber were present in *dy^3K^/dy^3K^* cross-sections, surrounded by myelinated nerves. In addition, Remak bundles in *dy^3K^/dy^3K^* that contain small axons (≤1.5 μm diameter) were not enveloped (as usually seen) or minimally enveloped in axonal groups. The naked larger and small axons in the sciatic nerve were similar to that observed in *dy^2J^/dy^2J^* [14]. AAV-CBh-αLNNdΔG2’ alone, or co-expressed with miniagrin only in muscle (SPc5-12 promoter), was sufficient to prevent amyelination of larger axonal bundles and well as with substantial increases of envelopment in Remak bundles. Double treatment similarly prevented amyelination of larger axons and Remak changes.

**Fig. 8.**
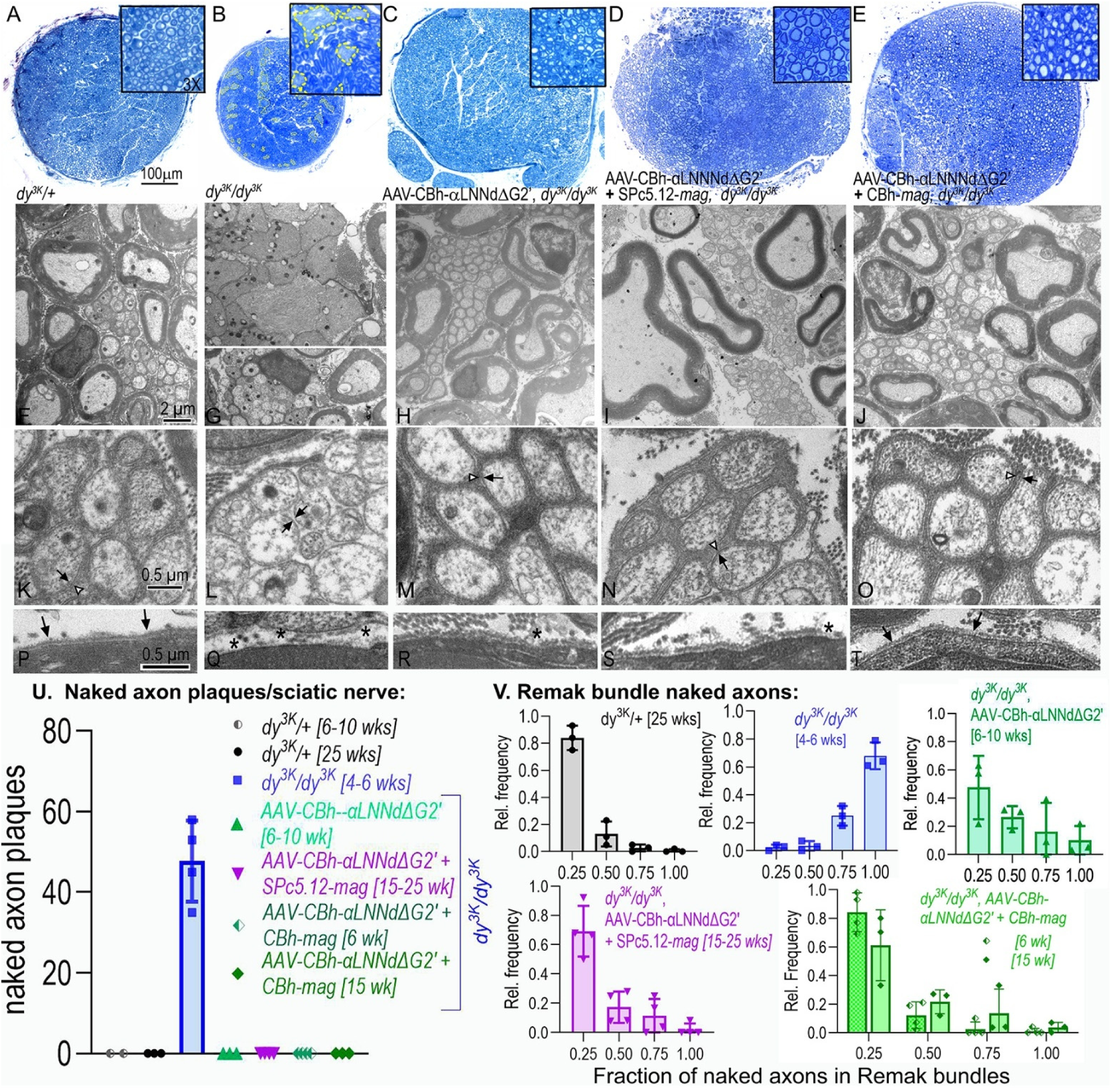
Sciatic nerve in dy3K. **A-E:** Methylene-blue (MeBlue) stained semithin sections for the indicated mice and treatments. Plaques of large adjacent naked axons (yellow outlined) were present in untreated *dy^3K^/dy^3K^*. **F-J, K-O:** *Remak bundle* axons (shown at higher magnification) are mostly enveloped in *dy^3K^/+* sciatic nerve (each axon membrane surrounded by a Schwann cell-derived membrane) while axonal membranes in untreated *dy^3K^/dy^3K^* lack an outer Schwann enveloping membrane. All AAV-treated Remak bundles are similar to *dy^3K^/+* with substantial axonal envelopment. **P-T***, Endoneurial BMs* shown at bottom for each condition. BMs (arrows) are thicker and more continuous with double AAV treatment (*, BM attenuations). **U**: Graph of count of naked axon plaques (patches) in MeBlue-stained sections (av ± s.d.). Naked bundles were not observed in any of the treated *dy^3K^/dy^3K^*. **V:** *Dy^3K^/dy^3K^* sciatic nerves also display naked axons (≤1.5 μm diameter) in Remak bundles. Normal levels of envelopment of these small axons were present following any of the AAV treatments. Frequency histograms of fraction of axons with 0-0.25, 0.25-0.5, 0.5-0.75, and >0.75 (av ± s.d.).

Given expression of both Lm411 and Lm421 detected in nerve, and given that β1 integrins are required for Schwann cell myelination [20, 26], we asked which compensating laminin might play a dominant role in myelination. Lm411 was previously found to bind poorly to β1-integrins [20, 27]. Is the same true for Lm421? This was addressed with a Schwann cell adhesion assay where it was found that Lm421, unlike Lm411, mediates β1-integrin dependent adhesion at concentrations comparable to known integrin-binding laminins (**Supplement Fig. 8**). In addition to integrins and dystroglycan, Schwann cell surfaces are enriched in sulfatides that bind to laminins through the α-subunit LG domains, enhancing laminin-mediated cell adhesion [28–30]. We evaluated an α4-laminin (Lm411) and found that it bound sulfatide with similar affinity compared to laminins 111 and 211.

We also evaluated linker-protein rescue of myelination by linker proteins through Lm411 and Lm421 in mouse dorsal root ganglia (DRG) organ cultures following isolation from LmC1^fl/fl^ pregnant mice [31]. After blocking endogenous laminin γ1 synthesis with adeno-cre, the DRGs were treated with α4-laminins without or with αLNNd/αLNNdΔG2’ linker proteins in the presence of ascorbate (to enable expression of collagen-IV also needed for myelination) (**Supplement Fig. 9**). Only limited myelination was detected with Lm411 + αLNNd while Lm421 +αLNNd generated more robust myelination. On the other hand, the combination of αLNNdΔG2’ plus miniagrin attached to Lm411 generated myelination equivalent to enablement of polymerization alone with Lm421. It is thought that the increased anchorage through miniagrin may indirectly enhance integrin binding and signaling.

Another question we addressed is whether direct laminin-integrin binding accounts for the totality of the myelination dependent upon laminin interactions. When Lm111 bearing a γ1E:Q mutation near the C-terminus, shown to be critical for laminin-integrin binding [32], was incubated with laminin-deficient DRGs, a significant reduction in myelination was observed (Suppl. Fig. 10, G-H). However, myelination was not completely abolished (as compared to absence of laminin). These results support previous findings using laminins 111 and 211 bearing a Lmγ1-C-terminal Flag tag found to impede integrin interactions [31]. Since published studies have revealed that polymerizing Lm111 and Lm211 can mediate BM assembly with recruitment of other integrin-binding components but *without* direct laminin-β1 integrin binding-activity [31, 33], we propose that the surviving myelinating activity depends only indirectly on laminins through its role in promoting BM assembly, i.e. integrin binding to the collagen-IV, perlecan or other recruited component.

## DISCUSSION

The small “linker” proteins αLNNdΔG2’, αLNNd and miniagrin were designed to bind to the γ1 short arm and coiled-coil of defective or compensating laminins, enabling polymerization by providing an αLN domain in a synthetic short arm, and enabling cell-anchorage to α-dystroglycan through the agrin LG domains attached to the long arm through the agrin NtA domain [14, 16, 28, 34]. Three mouse models (*dy^2J^/dy^2J^*, *dy^W^/dy^W^*, *dy^3K^/dy^3K^*) were previously shown to benefit with respect to the muscle pathology following muscle-specific transgenic expression of αLNNd in *dy^2J^/dy^2J^* [35], and αLNNd + miniagrin in *dy^W^/dy^W^* and *dy^3K^/dy^3K^* [15, 16].

Muscle-specific treatment, however, failed to correct the overriding neuropathy that causes a gait abnormality with partial paresis, one that progresses with increasing age. AAV-driven expression of αLNNdΔG2’ in the *dy^2J^/dy^2J^* using a universal (CBh) promoter, in which the linker bound to non-polymerizing α2-laminin, prevented sciatic nerve amyelination [14]. This was our first indication that a universal promoter could be used to drive expression of αLNNdΔG2’ in mice without undue or toxic effects.

We asked in this study whether AAV delivery of minigenes coding for αLNNdΔG2’ and miniagrin could be used to ameliorate the far more severe phenotypes of the *dy^3K^/dy^3K^* mouse in which there is a total absence of α2-laminins, with a particular focus on the contribution of laminin polymerization. The histological changes in muscle tissues and ultrastructural changes in sciatic nerve reflected improvements in survival, weights, strength and ambulation. The overall, increases in laminin subunits and receptors, as detected by immunofluorescence in muscle sarcolemma, were similar to those observed with muscle-specific transgenic expression of αLNNd and miniagrin in *dy^3K^/dy^3K^* mice [15]. Peripheral nerve branches, on the other hand, revealed some differences.

Sciatic nerve endoneurial Schwann cell (SC) BMs share the overall architectural organization with muscle sarcolemmal BMs; however, they differ in several respects: (i) β1-integrins, in particular α6β1 and α7β1 for laminins, are key mediators of myelination [19, 26] whereas in sarcolemma αDG is the paramount anchoring receptor [36]. (ii) While several SC integrins are available to interact with BM ligands, only one, α7β1_D_, appears to operate in mature muscle sarcolemma, with contribution of additional integrins (e.g. αv, α3) during regenerative myogenesis [37]. These other integrins interact with other BM components [26, 31]; however, matrix deposition of the BM proteins that provide other ligands requires laminin assembly [31]. (iii) While the α7 integrin knockout exhibits a mild phenotype in muscle and nerve, α7 integrin expression promotes regeneration and myelination [26, 38]. (iv) Lmα4, a normal SC subunit, contributes to myelination [17]. (v) SCs express gal-sulfatides [29] that increase α4-laminin cell adhesion. In Lmγ1 -/- dorsal root ganglion (DRG) cultures treated with recombinant laminins, ablation of laminin polymerization prevented radial axonal sorting and myelination [31].

Universal expression of αLNNdΔG2’ was striking in its ability to prevent the partial sciatic amyelination characteristic of the dy^3K^/dy^3K^ mouse, a predicted outcome. However, the absence of more than small increases of laminin subunits in *dy^3K^/dy^3K^* endoneurium was unexpected. In particular, this finding was different from the substantial endoneurial laminin increases that were seen with non-polymerizing α2-laminin following treatment of *dy^2J^/dy^2J^* mice with AAV-CBh-αLNNdΔG2’ [14]. The endoneurial findings in *dy^3K^/dy^3K^* mice suggest that the neural benefit did not derive from increasing Lm411 or Lm421 levels per se but rather benefitted from a structural reorganization of the BMs, likely through generation of laminin polymer stiffness [39]. It was shown in one recent study that SCs express a laminin- and collagen-IV binding G-protein receptor, Gpr126, that is needed for radial axonal sorting and myelination. Gpr126 has been found to be a mechanoreceptor whose activation requires laminin polymerization. Binding and polymer activation was found to be followed by increases of cAMP [40–42]. While Gpr126 was identified as binding to Lm211, our findings of Lm111 and polymer-enabled Lm411 myelination rescue imply that Gpr126 binds to several laminins [31, 43, 44]. Whether Gpr126 plays a distinct role when only interacting with collagen-IV, the other BM polymer, is not known. Schwann cell endoneurial BMs normally contain a mixture of α2- and α4-laminins, each contributing to radial axonal sorting and myelination with absence of each causing a partial loss of myelination and envelopment and absence of both causing severe amyelination as seen in nerve roots [17]. Given earlier measurements showing only partial occupancy of laminins achieved by AAV-αLNNdΔG2’ treatment, we strongly suspect expect that only a fraction of the α4-laminins was converted to a polymerizing form [14] with the non-polymerizing fraction providing the complementing laminin activity. Dystroglycan also plays a role in myelination, contributing to the stabilizing the myelin sheath and promotion of nodal development [25, 45]. Its primary contribution is thought to be that of cell anchorage to the cytoskeleton, although polymerization might be indirectly involved. A study by Xiao and colleagues [46] found that AAV-CB-*mag* treatment of the *dy^W^/dy^W^* hypomorphic mouse resulted in limited amelioration of nerve morphology, but did not prevent hindlimb paralysis and contractures.

The degree of underlying severity and linker protein-mediated phenotypic amelioration seen is less with *dy^3K^/dy^3K^* compared to *dy^2J^/dy^2J^*. One concern with *dy^3K^/dy^3K^* AAV treatments is that we had previously found the muscle tissue had very low (∼1/3) total γ1-laminin expression, reducing the laminins available for linker protein binding [15]. A recent study by colleagues examined the effect of dual universal transgene (CAG targeted to the Rosa26 locus) expression of αLNNd and miniagrin at presumed high expression levels in *dy^W^/dy^W^* and *dy^3K^/dy^3K^* and found substantial improvements in ambulation and grip strength with correction of the neuropathy [44]. The study increased the expectation that laminin α4 levels with sufficient (and perhaps early) linker expression would not be unduly limiting and served as a “gold-standard” for comparison. In the current study employing AAV coding for similar (αLNNdΔG2’ and *mag*) proteins, we saw comparable improvements in ambulation with, however, about two-fold lower weights and specific grip-strengths. Given that the BM alteration is inherently structural and stoichiometric in nature, we suspect the differences reflect lower and later expression using AAV as a minigene delivery system.

There are growing concerns that AAV doses need to be low for human therapy. Most AAV serotypes strongly transduce liver. Clinical trials have revealed that AAV-treated patients often develop hepatoxicity (with elevated serum ala/asp transaminases), occasionally leading to acute liver injury [47]. The frequency of some adverse events is thought to depend on the AAV dose (especially hepatotoxicity), and currently treatments employ < 2 x 10^14^ vg/kg [48–50]. Toxicity may also depend, in part, on AAV purity and the level of empty capsids.

The AAV doses used in the current study (as high as 5.8 x 10^14^ vg/kg) exceed the recommended doses for humans. The very high expression of αDG in muscle following treatment suggests that the dose of AAV-mag can be lowered. A way to further reduce dosage without loss of protein expression is to use a myotropic AAV, such as the two engineered AAV viruses shown to provide high muscle with minimal liver expression [51–53]. However, in the absence of an AAV to provide Schwann cell-specific expression αLNNdΔG2’ still needs a virus using a universal promoter. Further studies are needed to optimize delivery, balancing benefit against toxicity [54]. Of the mouse models, the *dy^3K^/dy^3K^* is the most challenging in this regard.

## METHODS

### AAV constructs

The construction of AAV_9_ containing mouse αLNNdΔG2’ and incorporating internal repeats (ITR), CBh promoter, full woodchuck response element (WPRE, fwp) and synthetic polyadenylation A signal (pA) was previously described [14]. This plasmid was modified by Vectorbuilder, Inc. (Chicago, IL) with a synthesized truncation of the WPRE element to remove a potentially oncogenic X-protein [55] and contain a short 49 nucleotide pA signal [56]. The mouse miniagrin viral sequence was provided as a gift by the Reinhard and Rüegg [44]. The DNA construct was synthesized and further modified by Vectorbuilder to replace the muscle specific promotor, SPc5-12, with a ubiquitously expressed CBh promoter, shortening of the full WPRE (swp) with the truncated version and swap of the bovine growth hormone pA with the synthetic version to enable efficient AAV encapsulation. All viral preparations were prepared by Charles River Laboratories (Kingston, NY).

### Mice and genotyping

**(a)** *Mice carrying the dy^3^****^K^*** allele, maintained in a mixed C57Bl/6/ 129SvEvTac background [15], were genotyped at the time of weaning or at post-natal day zero to one. Genomic DNA was purified from mouse tail or toe clipping respectively as previously described [15].

***Ambulation*** was recorded with VersaMax software (v 4.0) in 10 minute segments in an open-field box fitted with two rows of orthogonally placed light beam-block detectors as previously described [15].

### Microscopy and Morphometry

Forelimbs and hindlimbs were dissected from euthanized mice and processed as described [31]. (**a**) For paraffin embedding, muscles, following fixation as previously described [15], were sectioned at 5 μm, and stained with periodic acid Schiff (PAS) and Picro-Sirius Red (PSR) at the Histopathology Core Pathology Services (Rutgers University). Panoramic images were recorded with a Leica Digital Pathology Slide Scanner and analyzed with Aperio ImageScope software (Leica Biosystems, Nussloch, Germany). (**b**) For frozen sections, unfixed muscles were embedded in OCT, flash frozen in liquid nitrogen, followed by 5-μm sectioning with a cryostat at −20°C as described [15]. Sections were then washed, fixed in 3.2 % paraformaldehyde for 15 minutes at room temperature, and blocked in 5% goat serum overnight at 4°C[15]. Primary antibodies and secondary antibodies were incubated as described [15], followed by mounting with coverslips in 6% DABCO (1,4-diazabicyclo[2.2.2]octane) in glycerol. Detection of bound primary antibodies in fixed frozen sections was accomplished with Alexa Fluor 488 and 647 goat anti-rabbit, chicken, and mouse IgG secondary antibodies (Molecular Probes, Eugene, OR) at a 1:100 dilution. Tissue sections were stained together with antibodies concentrations to reveal differences in basement membrane component levels following different treatments. Regions of muscle were matched between genotypes, and the same exposure times and normalizations were applied to all images being compared.

### Antibodies

The following antibodies were employed at the concentration/dilutions as previously described [15] with the following specificities: polyclonal αLNNd specific antibody prepared against recombinant mouse αLN-LEa [15], rabbit antibody to mouse agrin (1:3000,kind gift from Markus Rüegg, Univ. Basel, Switzerland), 1 μg/ml chicken polyclonal antibody specific for the laminin-α4 subunit [12], 1 μg/ml chicken polyclonal laminin- α5 antibody [12], monoclonal rat anti-laminin β1 antibody (2 μg/ml, Invitrogen catalog # MAB14657), rat antimouse laminin γ1 antibody (Millipore catalog # 05-206, 1:100), rabbit anti-E4 (1μg/ml) [57], rabbit polyclonal anti-mouse laminin-β2 antisera, 1:3000 dilution [58, 59], rabbit anti-laminin β1/γ1 (0.5ug/ml, from anti-EHS [57], crossed-absorbed against immobilized recombinant laminin α1 short arm. Antibodies to detect receptors were rat anti-mouse β1 integrin (7.5 μg/ml, Millipore catalog # MAB1997), mouse anti-α-dystroglycan (1:100, Millipore catalog # 05-298), rat-anti mouse α7 integrin (5 μg/ml R & D cat# 33498). Antibody incubations to detect receptors have the addition of 0.1% Triton x-100 in the incubation mixture. Additional antibodies were rabbit anti-mouse collagen type IV (1:100, Millipore catalog # AB756P) and F4/80 monoclonal rat anti-mouse antibody (1:100; Abcam, catalog # ab6640) used to detect macrophages in frozen sections. Sections were counterstained with rabbit antiperlecan antibody (0.5 μg/ml; [40] or chicken anti-a4 laminin (1 μg/ml). Chicken antineurofilament-200 (1:1000, Abcam catalog# ab4680) and rat anti-mouse myelin basic protein (10 μg/ml;, Millipore AB9348) were used in dorsal root ganglia assays (31). Hamster β1-integrin-specific anti-rat IgM antibody Ha2/5 (BD Pharmingen, San Diego, CA, catalog # 555003) was used to block cell adhesion.

### Immunoblots

Mouse leg muscle at 15 weeks (40 mg) was pulverized in liquid nitrogen and lysed in 1ml buffer (50mM Tris pH 7.2, 150mM sodium chloride, 1mM EDTA, 1% SDS, 1% Triton x-100, protease inhibitors [Sigma catalog # P1860]) on ice for 20 min. Following centrifugation (15,000 x g), 20 ul of lysate was boiled in 1x Laemmli reducing buffer (with 5 % β-mercaptoethanol). Each sample was loaded (10 μl/lane) on an 8% SDS-polyacrylamide gel and transferred to PVDF membrane. Proteins were detected with rabbit anti-α1LN-LE (1 μg/ml), rabbit anti-mouse agrin (1:2500), or anti-GAPDH-HRP (1/35,000, Sigma catalog # G9295).

### Recombinant proteins

DNA constructs (with C-terminal (C) and N-terminal FLAG (*f*), c-Myc (*m*) in α1) for human Lmγ1, human Lmβ1(*Nha*), mouse Lmβ2 (*Nf*), mouse Lmα1 (*Nm*), mouse Lmα2 (*Nha*), mouse Lmα4 (*Nf*), mouse Lmα5 (*Cf*) were described previously [31, 33, 35, 60–62]. HEK293 cells, expressing Lm111, Lm411, Lm421, Lm511, and Lm521 heterotrimers tagged in the α or β subunit, were purified by antibody affinity chromatography for the tag contained in the secreted heterotrimer in conditioned medium were purified as described [28, 33]. Lm521 and Lm421 were generated by transfecting HEK293 cells previously stably expressing mouse β2 and human γ1 laminin under Zeocin G418 selection, respectively. This was followed by transfecting with either mouse Lmα5 or mouse Lmα4 DNA under puromycin selection. A glutamate to glutamine mutation (E:Q) was engineered into human γ1 laminin with overlapping PCR. A 5’ fragment (2528 nt) was generated with JABstXI EQ 1F 5’GCCAACCCTGTCCGTGTCCTGG3’ and JABstXI EQ 2R 5’CATCCAACGCAC- TGGCTAGGGCTTTTGAATG3’ as well as a 398 nt 3’ fragment using JABstX1 2F 5’CATTCAAAA- GCCCTAGCCAGTGCGTTGGATG3’ and JABStXI 1R 5’GATCTTCACGCGCCCTGTAGC3’. The 5’ and 3’ fragments were sewn together with the 1F and 1R primers (above) and digested with Bst X1 to produce a 2157 nt product that was ligated into the pCMV2 hg1 neo plasmid (Bst XI 9567 nt fragment, replacing the wildtype sequence). A stable trimeric laminin 111 γ1-E1607Q was made following transfection of a double-stable HEK293 cell line of mouse α1 laminin (N-terminal myc-tagged, puromycin) and human β1 laminin (N-terminal HA tagged, Zeocin) and cloning under triple antibiotics puromycin, zeocin and G418 (1 μg/ml, 100ug/ml, and 500 μg/ml respectively).

### Protein determinations

Recombinant laminin protein concentrations (molar) were determined from Coomassie blue-stained gel densitometry in comparison to an EHS-laminin (710 kDa protein mass) serving as standard ([38]) with molecular mass corrections for truncated laminins as previously described ([11, 39]). Absorbance at 280 nm was used to measure the concentration of αLNNdΔG2’ (111 kDa), αLNNd (156 kDa), and miniagrin (136 kDa).

### Dorsal root ganglia organ cultures

Dorsal root ganglia (DRGs) were dissected from E13.5 *Lamc1*fl^neo^/fl^neo^ embryos as described [31] and plated onto collagen-treated tissue culture wells with growth medium. The DRGs were treated 3 days later with purified cre-recombinase-GFP adenovirus. Recombinant nidogen-1 (28 nM), and recombinant laminins with/without laminin-binding linker protein (αLNNd or αLNNdΔG2’) were added with virus. After 3 additional days, the initial medium was replaced with myelination media containing ascorbic acid and the recombinant proteins. After 6 days the DRGs were fixed and prepared for immunostaining [31].

### Schwann cell adhesion assay

Costar 96-well half-area polystyrene dishes (no. 3695) were coated with BM proteins at the indicated concentrations in 0.15 M sodium bicarbonate (pH 9.6) overnight at 4°C. Plates were washed and blocked for 1 hr at RT with 0.3% heat-inactivated BSA in PBS. Rat Schwann cells, isolated from sciatic nerves (gift of Dr. James Salzer, New York University), were maintained in medium (passages 26-39) as described [63]. Cells were plated onto the coated wells (15,000 cells/well) and allowed to adhere for 70 min. Unbound cells were removed with three PBS washes. Schwann cells were fixed for 15 minutes at room temperature (RT, 10% acetic acid, 10% methanol) then wash 3x with 1x phosphate buffered saline. Crystal violet (5 mg/ml in 2% ethanol) was added to cells for 10 minutes at RT. The 96-well plate was submerged in a tank of water and rinsed 2 times followed by drying upside down. After adding 10% acetic acid for 10 minutes, the plates were read at 550nm on a Thermo Scientific Multiskan Sky plate reader.

### Sulfatide-binding assay

3Galß1-1’ceramide (brain galactosyl sulfatides) and galactosyl ceramide (Sigma C4905) were dissolved in methanol and 0.1 μg added per immulon-1B microtiter well. The plate was dried and the wells washed and blocked with ELISA Blocking Buffer (EBB; 1% BSA in TBS50/Ca). Proteins in EBB were added to each well and incubated for 1 hour at room temperature. Protein binding was detected with a HRP-linked monoclonal FLAG antibody (Sigma catalog # ab592) [28].

#### Statistics

Averages (Av) and standard deviations (s.d.) were calculated from measured values obtained from three or more images and morphometric measurements of perimeters and areas with the statistical packages in SigmaPlot 12.5, Excel, or GraphPad Prism (v.10). Averages and standard errors of the mean (s.e.m.) were determined from the means of consecutive sets of determinations (e.g. grip-strength) from different mice and from the means of myofiber cross-section areas from different mice. Three or more conditions were compared by one-way ANOVA followed by Holm-Sidak pairwise analysis in SigmaPlot (time-course plots) or Tukey’s multiple comparisons test in Prism (bar graphs). A difference was considered significant for P values < 0.05.

#### Study Approvals

The mouse protocol (9999-00384) and biosafety protocol (IBC# 13-574) for the study were approved by the respective IACUC and Institutional Biosafety Committees of Rutgers University – Robert Wood Johnson Medical School.

## Ackowledgements

This study was supported both by NIH grant R01-DK36425 (P.D.Y.) and NIH grant R01-AR085061 (P.D.Y.). Additional funding was provided by SEAL Therapeutics (University of Basel, Switzerland) contracted through the Rutgers University office of Life-Science Licensing and Technology, and Rutgers Health Bridge Funding.

