## Supplemental Figures 1-10 for "Dual AAV amelioration of Lama2-null muscular dystrophy and neuropathy"

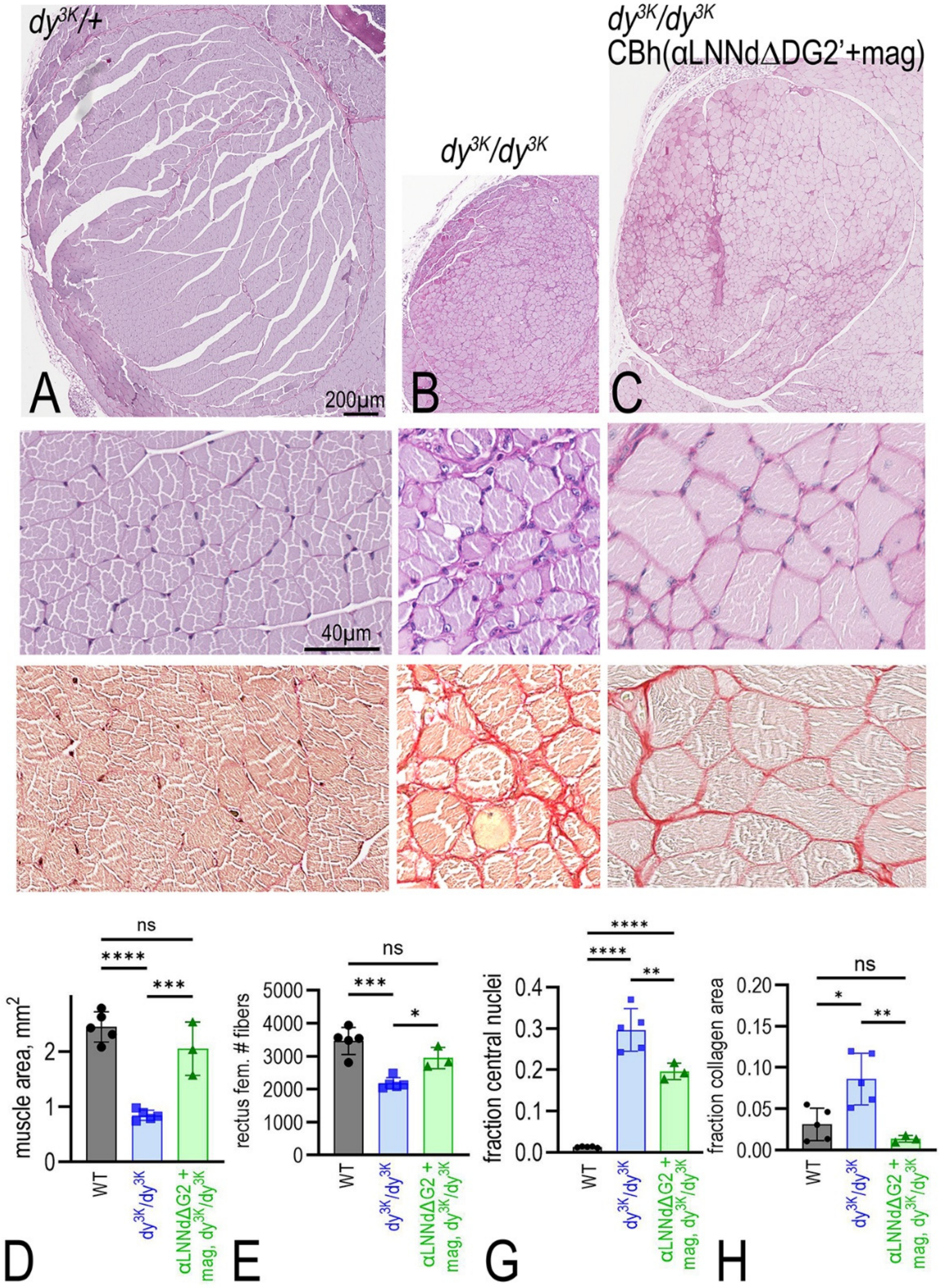


**Supplemental Fig. 1.** ***Histopathology of rectus femoris muscle.*** Hindlimb from wild-type, *dy*$^{3K}$/*dy*$^{3K}$, and *dy*$^{3K}$/*dy*$^{3K}$ treated with AAV-(CBh-αLNNdΔG2' + AAV-CBh-mag), were stained with periodic acid Schiff (PAS) and Picro-Sirius red (PSR). Treatment with AAV-(CBh-aLNNdΔG2' + CBh-mag) improved the morphology (angulation and regularity of myofibers, fewer central nuclei, less fibrosis) [**** P<0.0001; *** P<0.001; ** P<0.01; * P<0.05, 1-way Anova with Tukey's multiple comparisons test].

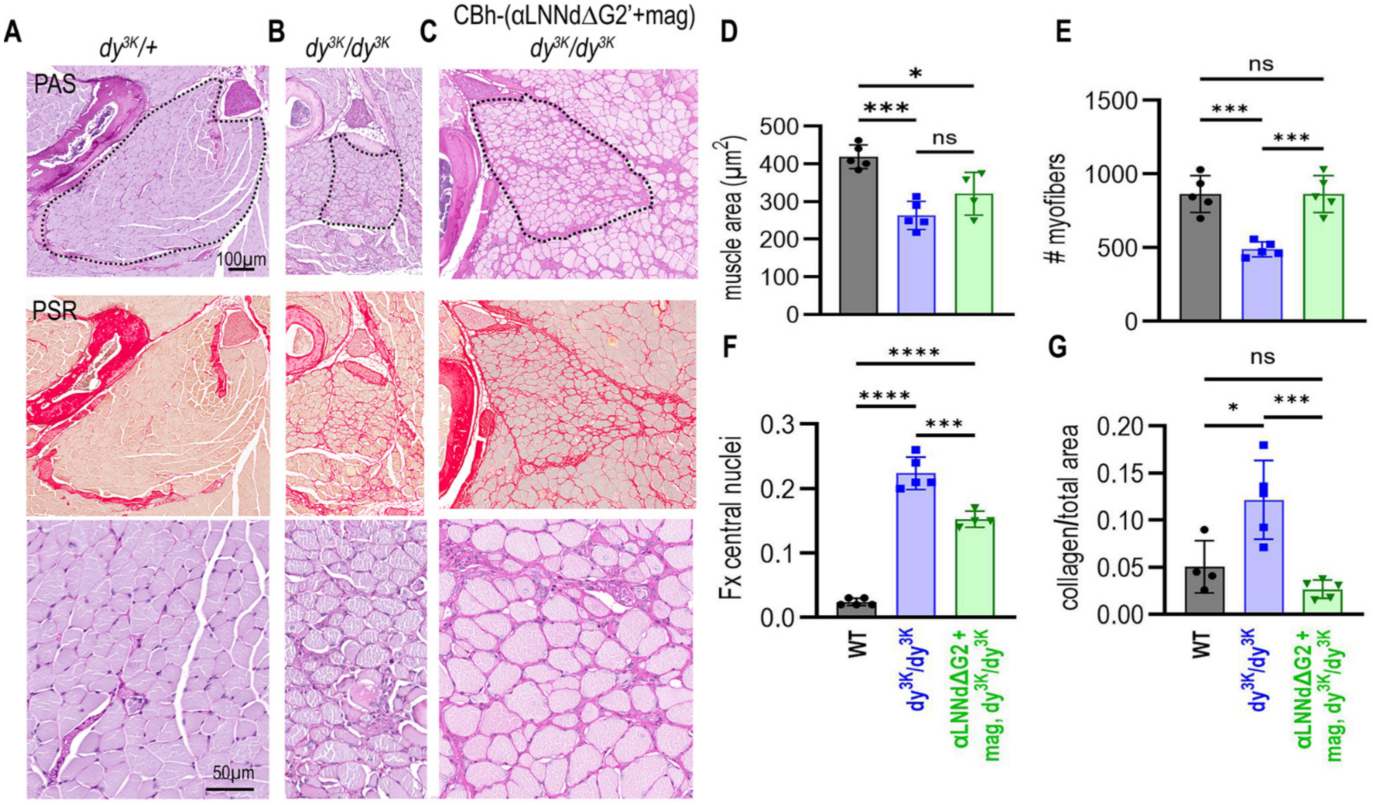


**Supplemental Fig. 2a**. *Histopathology of plantaris muscle.* Hindlimb from 6-week-old wild-type, *dy*$^{3K}$*/dy*$^{3K}$, and *dy*$^{3K}$*/dy*$^{3K}$ treated with AAV-(CBh-αLNNdΔG2' + AAV-CBh-*mag*), were stained with PAS and PSR. Treatment with AAV-(CBh-aLNNdΔG2' + CBh-mag) improved the morphology (angulation and regularity of myofibers, fewer central nuclei, less fibrosis). [**** $P<0.0001$; *** $P<0.001$; ** $P<0.01$; * $P<0.05$, 1-way Anova with Tukey's multiple comparisons test].

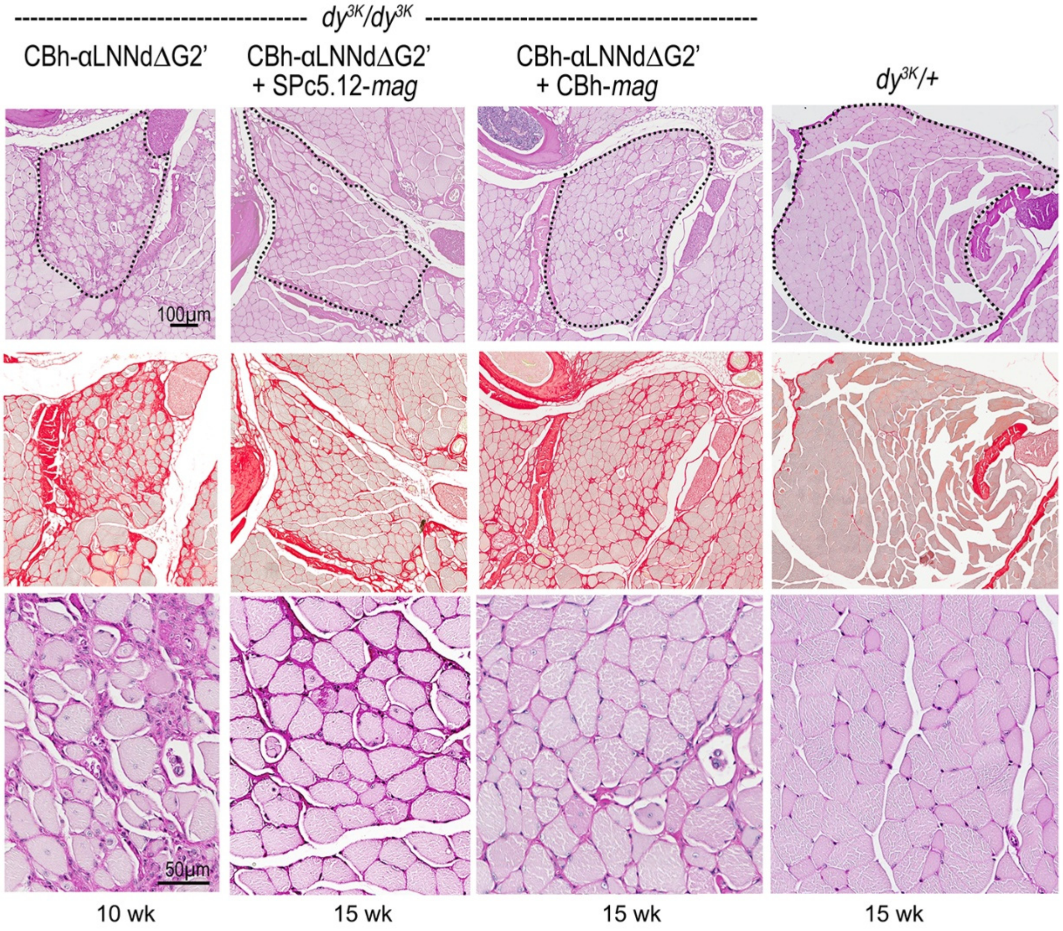


**Supplemental Fig. 2b**. *Histopathology of adult plantaris muscle.* Hindlimb from adult *dy*$^{3K}$/+ and *dy*$^{3K}$/*dy*$^{3K}$ treated with AAV$_9$-CBh-αLNNdΔG2', AAV$_9$-CBh-αLNNdΔG2' + AAV SPc 5-12-*mag*, or AAV$_9$-CBh-αLNNdΔG2' + (AAV$_9$-CBh-*mag*) were stained with PAS and PSR. Treatment with AAV$_9$-CBh-aLNNdΔG2' + AAV$_9$--SPc5-12 or AAV$_9$-CBh-*mag* improved the muscle morphology (angulation and regularity of myofibers, fewer central nuclei) compared to *dy*$^{3K}$/*dy*$^{3K}$ with a single AAV$_9$-CBh-αLNNdΔG2' treatment (there were no surviving *dy*$^{3K}$/*dy*$^{3K}$ after 8 weeks for comparison).

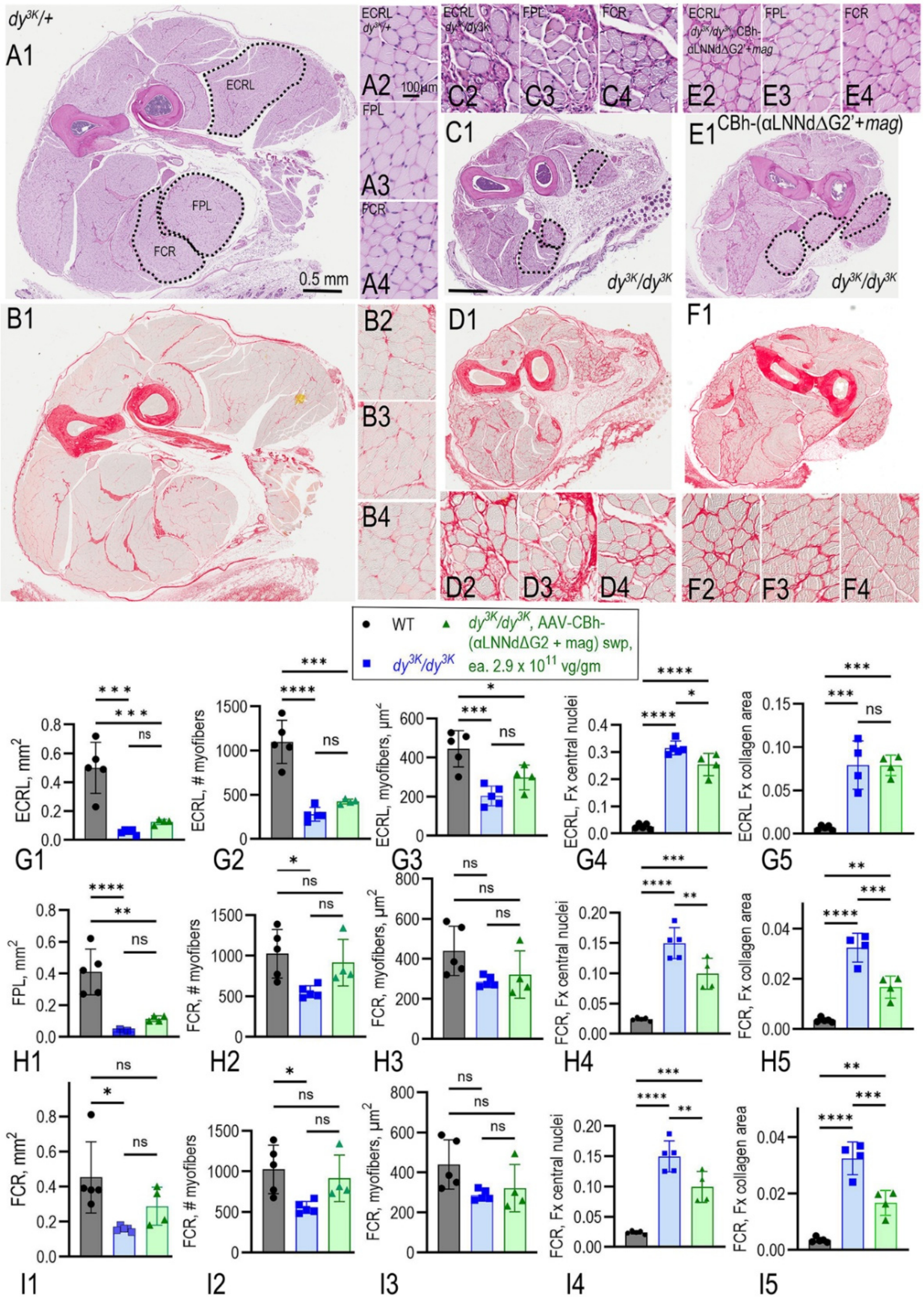


**Supplemental Fig. 3**. *Muscle histology of distal forelimb*. Cross-sections from 6-week-old mice were stained with PAS and PSR. Extensor carpi radialis longus (ECRL), flexor policis longus (FPL) and flexor carpi radialis (FCR), small muscles contributing to gripping, were examined to determine average overall size, fiber count, myofiber area, fraction (Fx) central nuclei, and fraction of PSR-positive (collagen) muscle. Reductions in central nuclei and collagen were observed in *dy*$^{3K}$/*dy*$^{3K}$ treated with αLNNdΔG2' + *mag* with little or no increase in muscle size and myofiber count. [**** P<0.0001; *** P<0.001; ** P<0.01; * P<0.05, 1-way Anova with Tukey's multiple comparisons test].

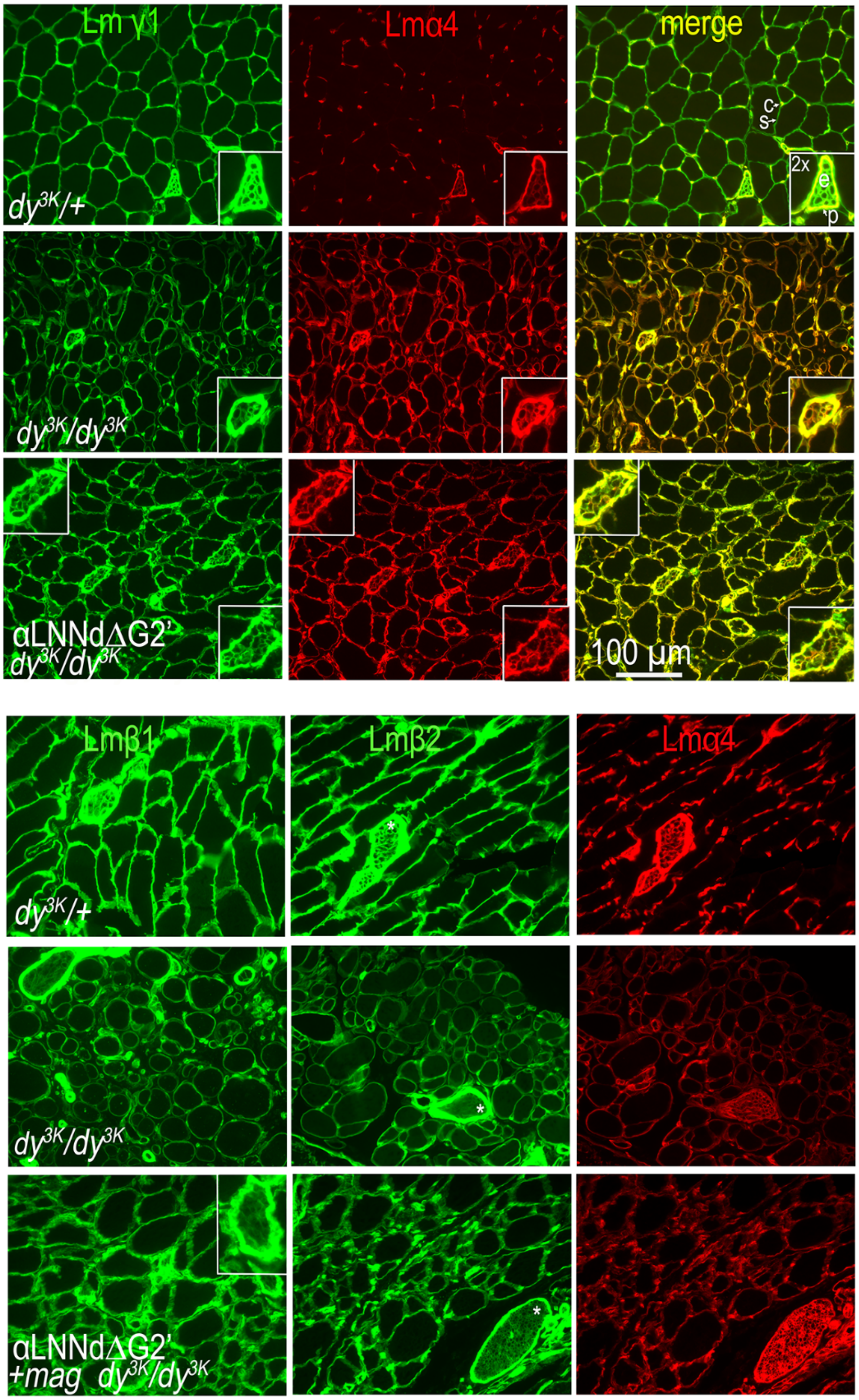


**Supplemental Fig. 4.** *Immunohistology, laminin subunits.* 6- week-old triceps muscle was stained with antibodies for Lmγ1, Lmα4, Lmβ1, and Lmβ2. Insets from the same or adjacent fields show nerves at 2x magnification. Sarcolemmal (s), capillary (c), endneurial (e) and perineurial (p) BMs indicated in upper right panel. The sarcolemmal BM, seen between capillaries, and endoneurial BMs of small nerve branches (* or inset), were reduced in intensity for Lmβ1, Lmβ2 and Lmγ1 subunits in $dy^{3K}/dy^{3K}$. AAV delivery of αLNNdΔG2' + *mag* in $dy^{3K}/dy^{3K}$ partially increased intensity of subunit staining in sarcolemmal BMs, but to levels below $dy^{3K}/+$. Endoneurial laminin intensities were modestly increased in αLNNdΔG2' + *mag* in $dy^{3K}/dy^{3K}$ in some but not all nerves relative to untreated $dy^{3K}/dy^{3K}$.

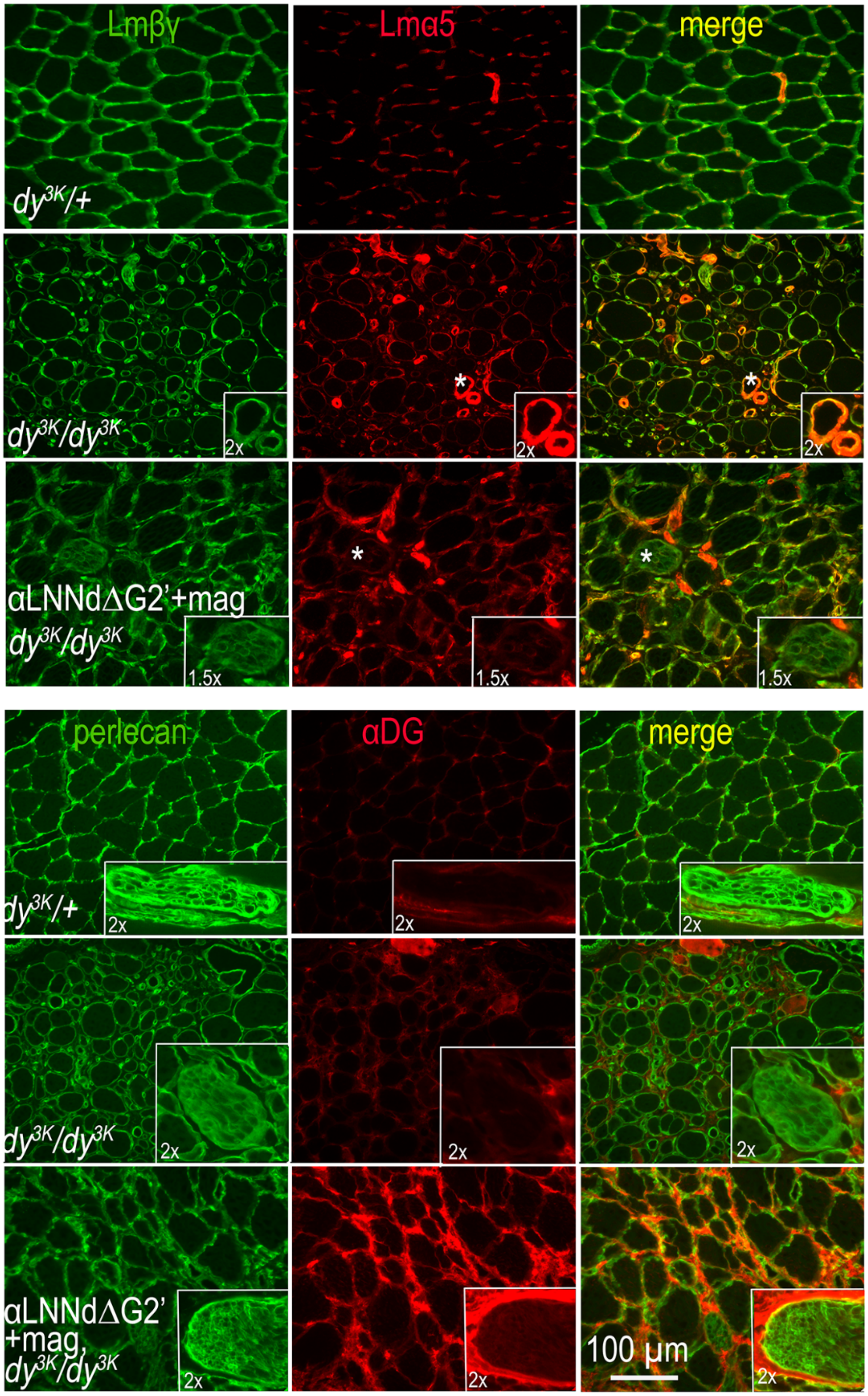


**Supplemental Fig. 5.** *Immunohistology, laminin-α5 and α-dystroglycan.* Triceps muscle (6 wks age) were stained with antibodies to laminin-α5 and α-dystroglycan (αDG. Lmα5 is normally detected in only muscle capillaries. In untreated and double-linker treated *dy*$^{3K}$/*dy*$^{3K}$, staining intensity is increased the microvasculature and detected in sarcolemma. αDG staining was modestly increased in the untreated dystrophic muscle and greatly increased with combined αLNNdΔG2' and *mag* expression. Nerve 2x inserts shown from other fields from the same muscle. Unlike sarcolemma and epineurium, little αDG was detected in cross-sections of muscle to show endoneurial BMs.

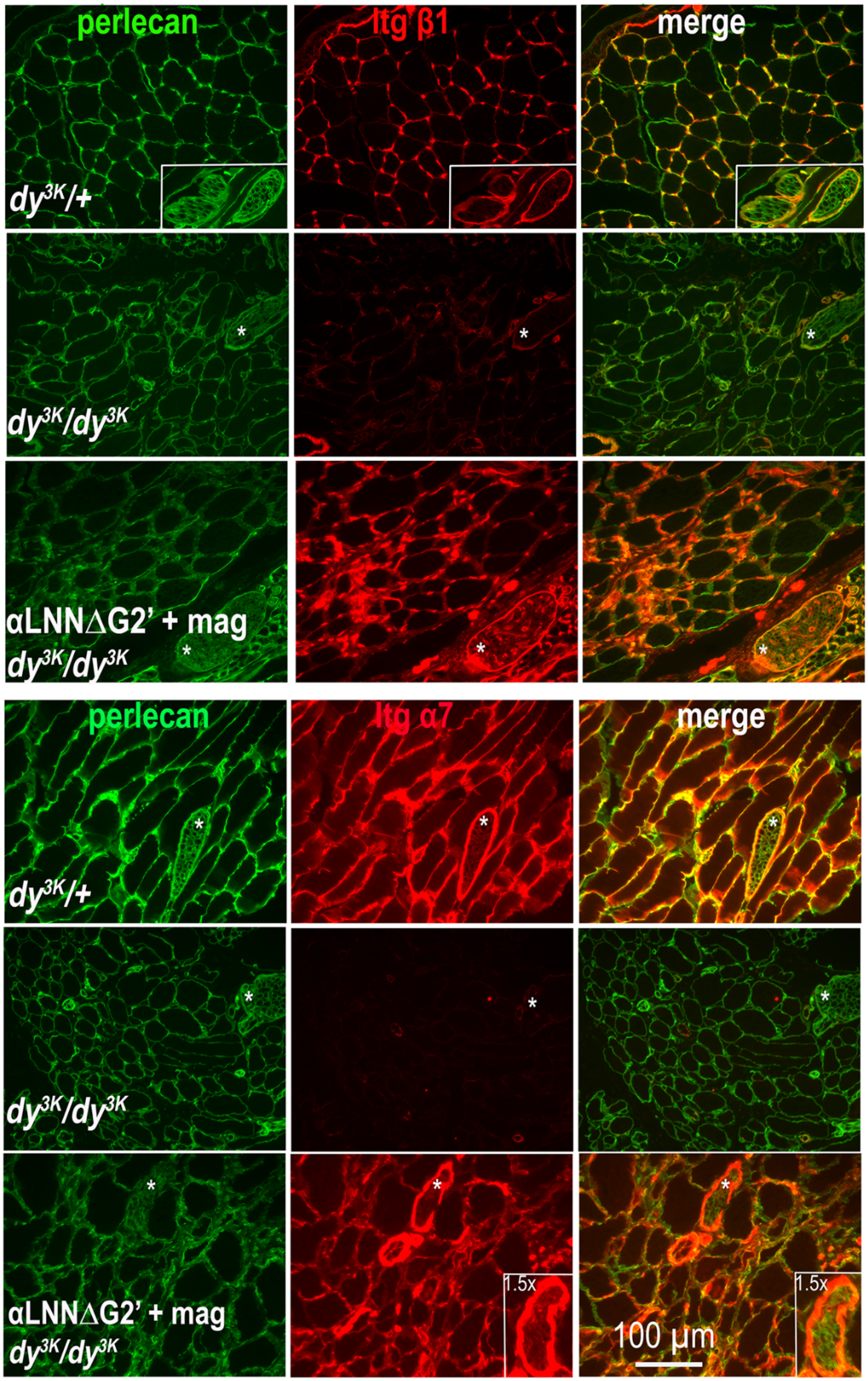


**Supplemental Fig. 6.** *Immunohistology, integrins.* Triceps muscle (6 weeks) was stained with monoclonal antibodies to integrins β1 and α7 shown compared to perlecan. Expression of αLNNdΔG2' and *mag* was associated with higher intensity levels of the two integrin subunits compared to untreated dystrophic mice.

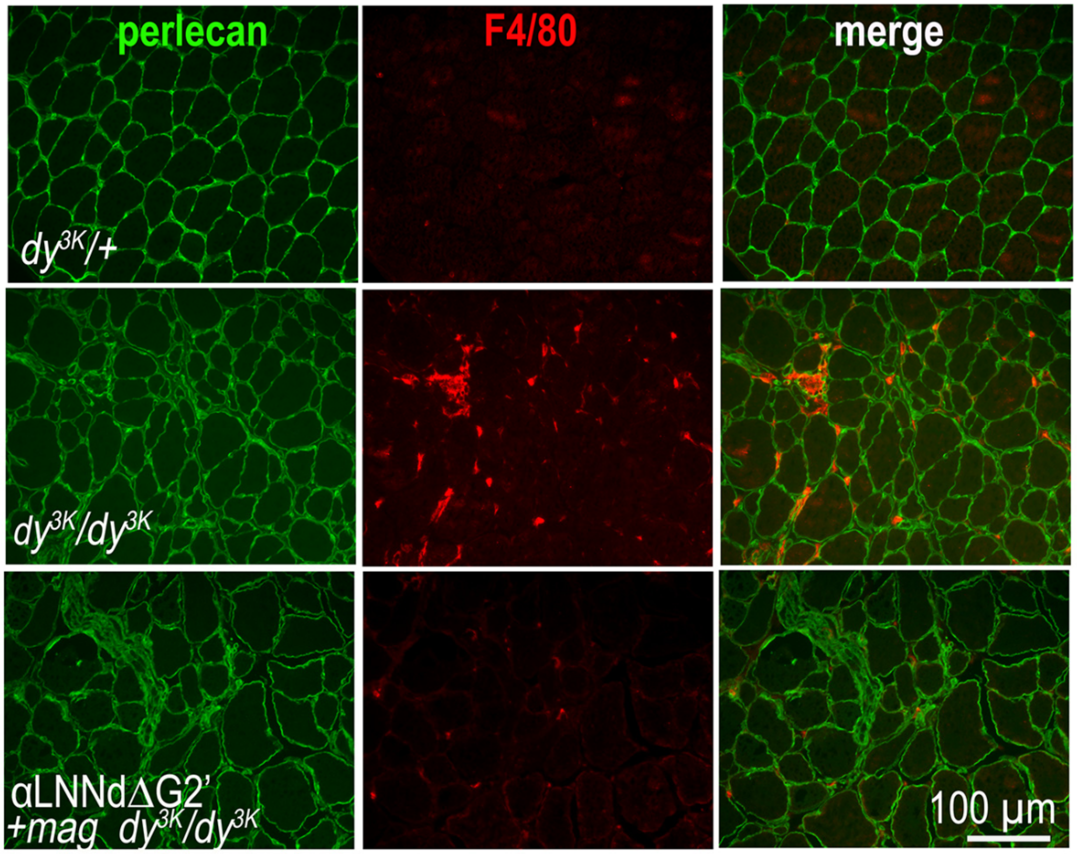


**Supplemental Fig. 7**. *Inflammation*. Triceps muscle (6 weeks age) was stained with the antibody F4/80 (red), a specific macrophage marker, and counterstained for perlecan, a BM marker (green). Macrophages, absent in *dy*$^{3K}$/+, were abundant in untreated *dy*$^{3K}$/*dy*$^{3K}$ and considerably reduced in *dy*$^{3K}$/*dy*$^{3K}$-αLNNdΔG2' + *mag*.

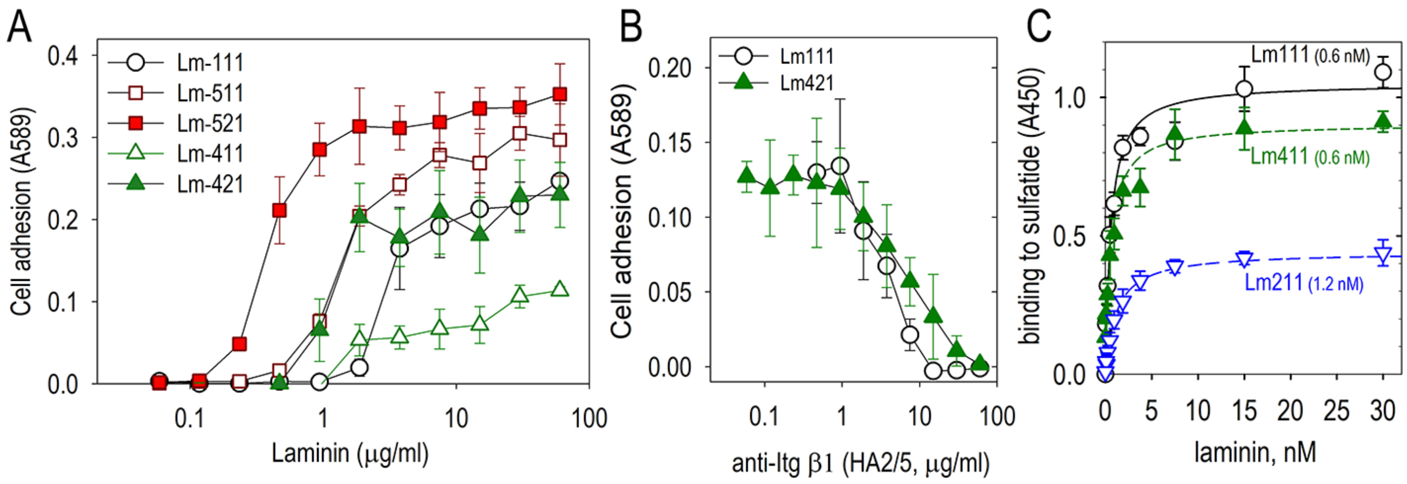


**Supplemental Fig. 8**. *Schwann cell adhesion to $\alpha$4-laminins and sulfatide binding.* **A**. Schwann cells (15,000/well) were incubated in 96-well half-bezel wells coated with the indicated recombinant laminins. **B**. Schwann cells were incubated with anti-integrin β1 antibody HA2/5 at the indicated concentrations. After washing, cells were stained with crystal violet with absorbance determined at 589 nm. Lm411 supported cell adhesion poorly compared to Lm421 and other laminins. Lm421 adhesion, like that of Lm111, was completely inhibited by the integrin β antibody. **C**. The indicted laminins were incubated in serial 2-fold dilutions on plates coated with galactosyl-sulfatide (n=3 replicates). Bound laminins were detected with anti-E4 (Lmβ1/β2 LN-LEa specificity) antibody.

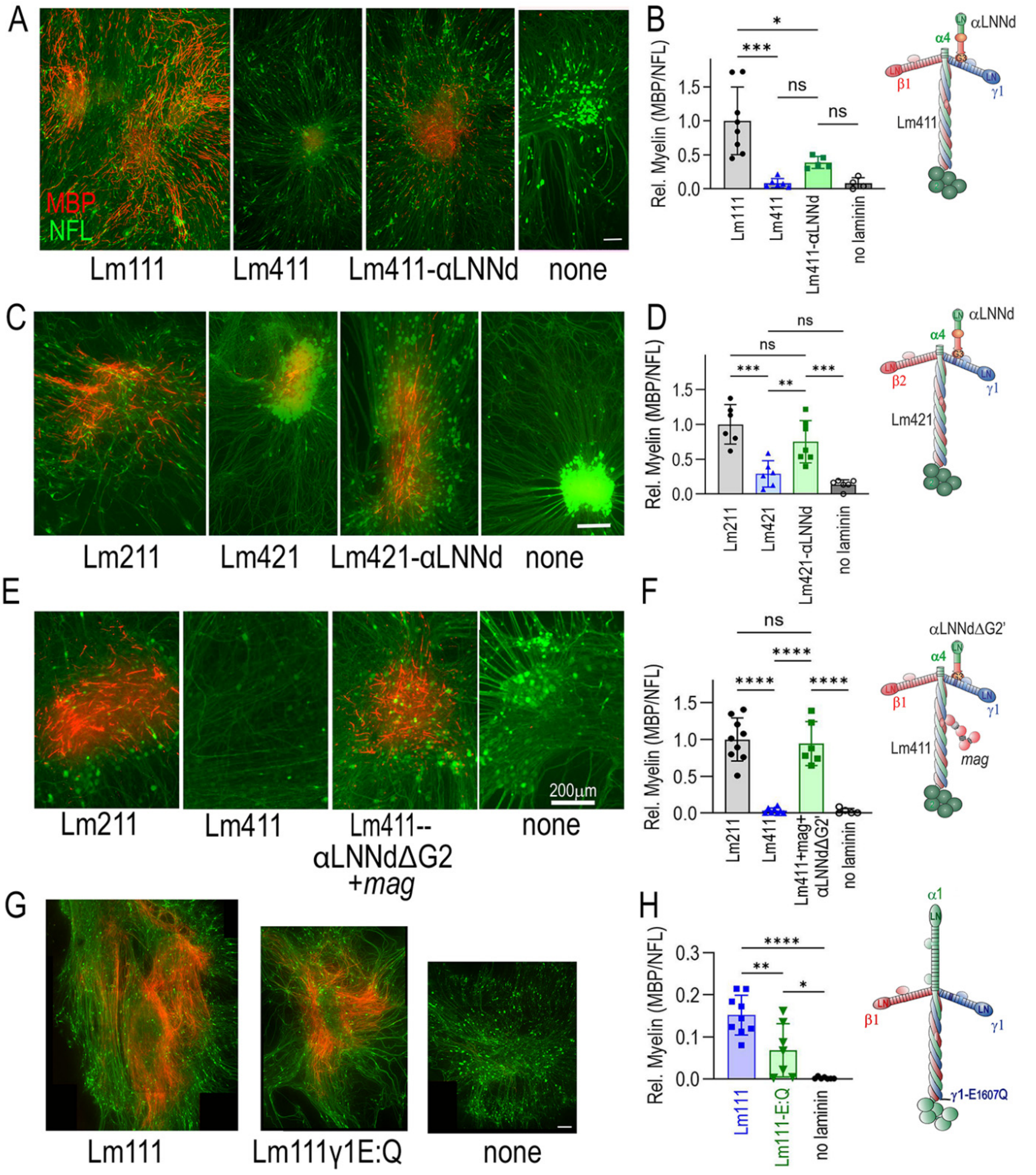


**Supplemental Fig. 9**. *Myelination in Lm$\gamma$1-/- dorsal root ganglia*. DRGs excised from pregnant E13.5 Lm$\gamma 1^{fl/fl}$ mice were grown in culture for 3 days and treated with cre-adenovirus to inactivate the LamC1 gene. The cultures were then treated with the indicated laminins with nidogen-1 in the presence of ascorbate without or with the indicated linker proteins. (**A,B**): Lm111 and Lm411 +/- linked to αLNNd + nidogen-1 (28 nM each); (**C,D**): Lm211 and Lm 421 +/- linked to αLNNd + nidogen-1 (28 nM each); (**E,F**): Lm211 and Lm411+/- linked to αLNNdΔG2' + nidogen-1 (14 nM each). Bar plots shown in B,D,F,G (average ± sd from DRGs in stitched 4x fields; **** P<0.0001; *** P<0.001; ** P<0.01; *P<0.05). Lm421 linked to αLNNd was sufficient to enable myelination of DRGs while apparent myelin increases with Lm411-αLNNd were not significant. (**G,H**): Lm111 with a γ1-E1607Q mutation (E1605Q in mouse) that prevents integrin binding to laminin, reduced but did not eliminate myelination. Length bars (200 μm). [**** P<0.0001; *** P<0.001; ** P<0.01; * P<0.05, 1-way Anova with Tukey's multiple comparisons test].

*dy*$^{3K}$/+ *v. dy*$^{3K}$/*dy*$^{3K}$ 6 wks | *dy*$^{3K}$/+ v. *dy*$^{3K}$/*dy*$^{3K}$ - CBh αLNNdΔG2', 10 wks | *dy*$^{3K}$/+ v. *dy*$^{3K}$/*dy*$^{3K}$ - CBh (αLNNdΔG2'+mag), 9.5 wks

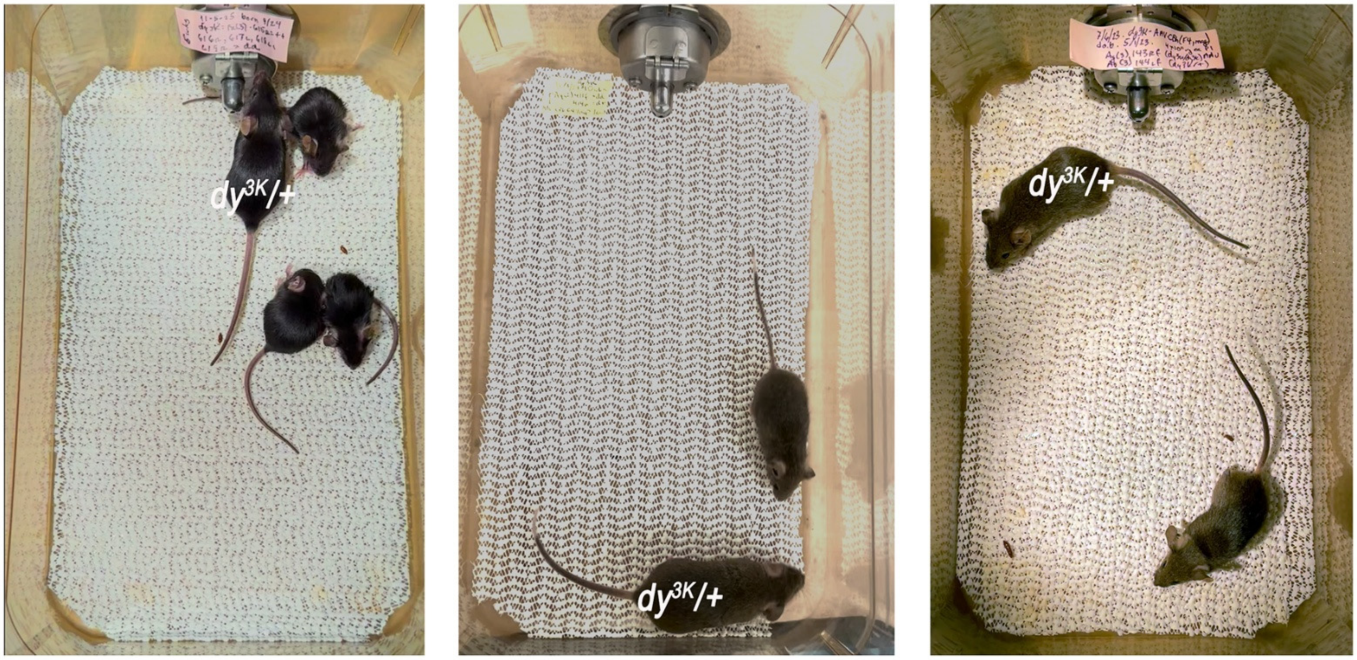


Video-1 | Video-2 | Video-3

**Supplemental Fig. 10**. *Videos of dy*$^{3K}$*/+ and dy*$^{3K}$*/dy*$^{3K}$ mice: Non-labeled mice are homozygous mutants.
